# RNA splicing variants of the novel long non-coding RNA, CyKILR, possess divergent biological functions in non-small cell lung cancer

**DOI:** 10.1101/2024.07.08.602494

**Authors:** Xiujie Xie, H. Patrick Macknight, Amy L. Lu, Charles E. Chalfant

**Author notes:** To whom correspondence should be addressed: Charles E. Chalfant, Professor, Department of Medicine, Division of Hematology & Oncology, P.O. Box 801398, University of Virginia-School of Medicine, Charlottesville, VA, 22903, or. Contributed equally as senior authors (Xiujie Xie, Charles E. Chalfant).

## Abstract

The CDKN2A gene, responsible for encoding the tumor suppressors p16(INK4A) and p14(ARF), is frequently inactivated in non-small cell lung cancer (NSCLC). Herein, an uncharacterized long non-coding RNA (lncRNA) (ENSG00000267053) on chromosome 19p13.12 was found to be overexpressed in NSCLC cells with an active, wild-type CDKN2A gene. This lncRNA, named **<u>Cy</u>**clin-Dependent **<u>K</u>**inase **<u>I</u>**nhibitor 2A-regulated **l**nc**<u>R</u>**NA (CyKILR), also correlated with an active WT STK11 gene, which encodes the tumor suppressor, Liver kinase B1. CyKILR displayed two splice variants, CyKILRa (exon 3 included) and CyKILRb (exon 3 excluded), which are cooperatively regulated by CDKN2A and STK11 as knockdown of both tumor suppressor genes was required to induce a significant loss of exon 3 inclusion in mature CyKILR RNA. CyKILRa localized to the nucleus, and its downregulation using antisense RNA oligonucleotides enhanced cellular proliferation, migration, clonogenic survival, and tumor incidence. In contrast, CyKILRb localized to the cytoplasm, and its downregulation using siRNA reduced cell proliferation, migration, clonogenic survival, and tumor incidence. Transcriptomics analyses revealed enhancement of apoptotic pathways with concomitant suppression of key cell cycle pathways by CyKILRa demonstrating its tumor-suppressive role. CyKILRb inhibited tumor suppressor microRNAs indicating an oncogenic nature. These findings elucidate the intricate roles of lncRNAs in cell signaling and tumorigenesis.

## INTRODUCTION

Non-small cell lung cancers (NSCLCs) encompass a diverse group of lung malignancies, accounting for approximately 85% of all lung cancer cases.[1–4] Despite significant progress in diagnosis and treatment, NSCLCs remain a leading cause of cancer-related deaths worldwide.[5] The intricate molecular landscape of NSCLCs, characterized by complex tumor microenvironments and diverse genetic alterations, presents considerable challenges in understanding their pathogenesis and developing effective therapeutic strategies. In regard to genetic alterations, the cyclin-dependent kinase inhibitor 2A (CDKN2A) gene encodes the tumor suppressors, p16(INK4A) and p14(ARF), and is found inactivated or deleted in NSCLC at approximately 30% [6, 7] . p14ARF is an alternate reading frame protein product of the CDKN2A locus, and p14ARF is induced in response to elevated mitogenic stimulation, such as aberrant growth signaling from the *MYC* and *RAS* genes. It accumulates mainly in the nucleolus where it forms stable complexes with NPM or Mdm2. p16INK4a (also known as cyclin-dependent kinase inhibitor 2A, multiple tumor suppressor 1, and numerous other synonyms) is a protein that suppresses cell division by slowing the progression of the cell cycle from the G1 phase to the S phase, thereby acting as a tumor suppressor. Loss of either p14ARF or p16INK4a has dramatic effects on the landscape of gene expression and RNA transcription increasing the susceptibility of cells to undergo oncogenic transformation.

Effects on gene expression can also include long noncoding RNAs (lncRNAs), which were for many years dismissed as “transcriptional noise”. In recent years, lncRNAs have gained significant attention due to their potential roles in various diseases including cancer.[8–11] LncRNAs are RNA transcripts with lengths exceeding 200 nucleotides, and differ from protein-coding genes in not encoding for functional proteins. Instead, they exert their biological effects through diverse molecular mechanisms. With unique secondary and tertiary structures, lncRNAs can regulate gene expression through processes like chromatin modification, RNA splicing, and mRNA transcriptional and post-transcriptional regulation via RNA-DNA, RNA-microRNA, or RNA-Protein interactions.[12–16] In cancer biology, dysregulation and aberrant expression of lncRNAs have been linked to tumor initiation, progression, and therapeutic responses.[17] Distinct lncRNA expression patterns have been observed in various cancer types, including lung, breast, prostate, and colorectal cancers, among others.[18–22] Furthermore, accumulating evidence suggests that lncRNAs can act as oncogenes or tumor suppressors exerting both pro-tumorigenic or anti-tumorigenic effects.[23, 24]

Like protein-coding genes, lncRNA genes consist of introns and exons that undergo alternative splicing. LncRNA splicing plays a pivotal role in generating transcriptomic diversity and expanding the functional repertoire of lncRNAs. Alternative splicing events give rise to distinct isoforms of lncRNA, which can differ in their sequences, structures, and subcellular localization. These splice variants possess the remarkable capacity to differentially modulate gene expression and cellular function in a context-dependent manner.[25–27] Although lncRNAs and their splicing variants have emerged as crucial players in the intricate landscape of cellular gene regulation,[28] identifying lncRNAs and their various splice variants remains challenging despite advancements in high-throughput sequencing technologies and bioinformatics approaches. Notably, one identified lncRNA, Neat1, undergoes alternative splicing regulated by RBM10. The resulting two splice variants, Neat1-1 and Neat1-2, play distinct roles in the invasion and metastasis of NSCLCs.[29] Another reported lncRNA has multiple splice variants, SOX2OT (4 & 7), which have oncogenic roles in NSCLCs.[30] Thus, the alternative splicing of lncRNAs is becoming appreciated for their distinct and novel roles in NSCLC and cancer, but are significantly understudied.

In this study, we have made the exciting discovery of a novel lncRNA (ENSG00000267053) located on the forward strand of chromosome 19p13.12 that exhibits multiple splice variants. This lncRNA, termed **<u>Cy</u>**clin-Dependent **<u>K</u>**inase **<u>I</u>**nhibitor 2A-regulated **l**nc**<u>R</u>**NA (CyKILR), is almost exclusively present in NSCLC cells with both an active and wild-type *CDKN2A* and *STK11* gene. Based on the presence of exon 3, the CyKILR splice variants are classified into two distinct groups, CyKILRa (exon 3 included) and CyKILRb (exon 3 excluded). Mechanistic studies show that CyKILRa predominantly localizes in the nucleus and demonstrates a tumor-suppressive function. In contrast, CyKILRb localizes to the cytoplasm and exhibits prosurvival functions. The balance between the two groups of CyKILR RNA splicing variants is dynamically regulated by gene products of both the *CDKN2A* and *STK11* genes, thereby finely tuning cellular homeostasis in these NSCLC cell lines. Our findings shed light on the potential diagnostic, prognostic, and therapeutic implications of targeting lncRNAs and their specific splicing variants in NSCLCs opening new avenues for precision medicine approaches. Indeed, this study also unravels the intricate regulatory networks orchestrated by lncRNAs and their splice variants offering an exciting opportunity to decipher their complex roles in cancer biology.

## RESULTS

### The association of the expression of a novel long non-coding RNA and CDKN2A/LKB1 status in NSCLCs

The *CDKN2A* gene is found inactivated or deleted in NSCLC at a high percentage, which also means that a number of NSCLC tumors and cell lines have a wild-type (WT), active, and functional *CDKN2A* gene. To examine if there are specific gene expression patterns between WT *CDKN2A* versus mutant/inactive *CDKN2A* NSCLC cell lines, twelve NSCLC cell line with a differing *CDKN2A* status were compared (six with WT *CDKN2A* (H838, H522, H2009, H1299, H1792, and H1734) and six with deleted/mutant *CDKN2A* (H2030, H810, H460, A549, H838, and H1648)) using unbiased transcriptomics. A top “hit” for a differentially expressed lncRNA associated with a WT *CDKN2A* gene was a novel, uncharacterized lncRNA (ENSG00000267053) (**Table S1**) located on chromosome 19p13.12 (**Figure S1**). Validation studies using qRT-PCR corroborated a strong link between this **<u>Cy</u>**clin-Dependent **<u>K</u>**inase **<u>I</u>**nhibitor 2A-regulated <u>**l**</u>nc**<u>R</u>**NA (CyKILR) and the WT *CDKN2A* gene, but a perfect correlation was not observed (**Fig.1A-C**). Indeed, the DESeq2 and plot3 packages in R studio were also utilized to generate both 3D principal component analysis (PCA) and correlation plots of CDKN2A mRNA expression with CyKILR expression, but the PCA showed that the 12 NSCLC lines did not exhibit strong clustering based solely on CDKN2A status/expression (**Fig.1D**). Examining additional oncogenotypes to explain the inconsistency showed that the five NSCLC cell lines with high CyKILR expression (H358, H522, H2009, H1299, H1792) also correlated with the expression of the tumor suppressor, Liver kinase B1 (LKB1), a product of the *STK11* gene (**Fig.1E,F**). The protein levels of the *CDKN2A* gene products, p14 ARF and p16 INK4A, as well as LKB1 were also examined to validate our transcriptomic and qRT-PCR findings (**Fig.1G,H**). These results were consistent with the mRNA expression findings across the 12 cell lines. Importantly, our analyses revealed high expression of CyKILR in only H358, H522, H2009, H1299, and H1792, which were cell lines positive for both WT *CDKN2A* and *STK11* genes and demonstrated expression of their gene products at the protein level (**Fig.1G,H)**. These results demonstrate that in NSCLC cell lines, CyKILR is almost exclusively expressed when the cells possess both active and WT *CDKN2A* and *STK11* genes.

**Figure 1.**
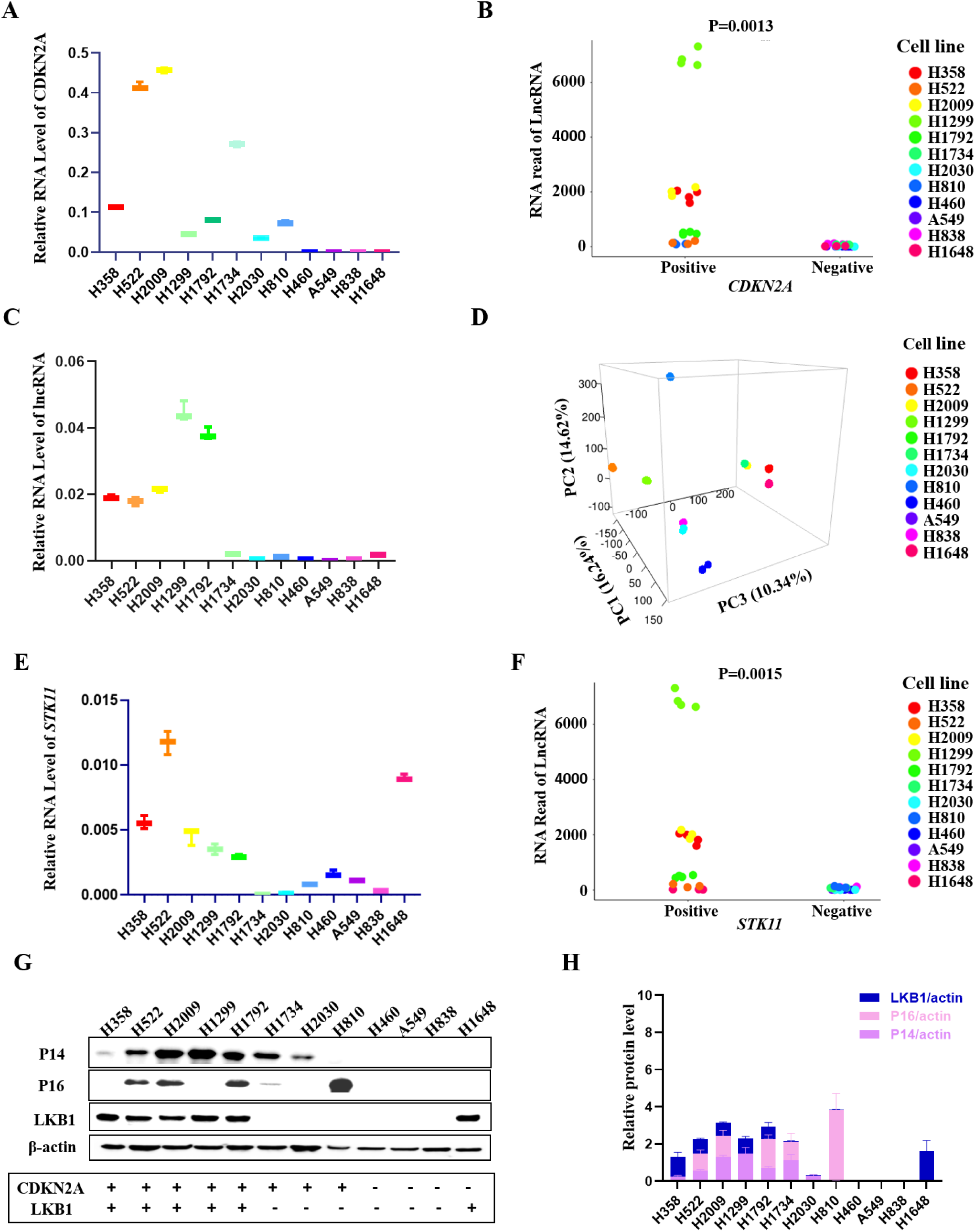
The association of CyKILR expression with an active, wild-type *STK11* and *CDKN2A* genes in NSCLCs. Total RNA was collected from the indicated NSCLC cell lines and reverse transcribed (RT). Quantitative Real-Time PCR (qRT-PCR) and deep RNA sequencing were conducted. The R program was used for analysis of deep RNA sequencing data. Western immunoblotting was conducted on 12 NSCLC lines to assess the expression of *CDKN2A* and *STK11* gene products with CyKILR expression. **A.** The mRNA level of CDKN2A across the 12 NSCLC lines normalized to GAPDH mRNA levels. **B.** Correlation plot of deep RNA sequencing data displaying the relationship between *CDKN2A* gene status and the expression of the lncRNA, CyKILR. **C.** CyKILR RNA levels by qRT-PCR across the 12 NSCLC lines normalized to GAPDH mRNA levels. Data in Panels A and C are presented as mean□(SD) (n□≥□3 replicates). **D.** Three-dimensional PCA plot representing the variation of CDKN2A and CyKILR expression for the 12 NSCLC lines. **E.** The mRNA level of the *STK11* gene product, LKB1, across the 12 NSCLC lines normalized to GAPDH mRNA levels. Data in Panel E are presented as mean□±□SD (n□≥□3 replicates). **F.** Correlation plot of deep RNA sequencing data to an active, wild-type *STK11* gene displaying the relationship between *STK11* mRNA expression and CyKILR expression. **G.** Relative protein levels of the indicated factors across a panel of 12 cell lines with β-actin used for normalization. Data are representative of n=3. **H.** The protein levels of the *CDKN2A* gene products, p14 and p16, the *STK11* gene product, LKB1, across the 12 NSCLC lines with β-actin used for normalization. Data in Panel H are presented as mean□±□SD (n□≥□3 replicates). Data in Panels B and F were analyzed by unpaired t-test with Welch’s correction. Reverse transcription (RT) “minus” controls (-RT) were included in all RT-PCR experiments and replicates with no signal observed.

### CyKILR classifies into two distinct isoform groups based on the alternative RNA splicing of Exon 3

RNA sequencing data revealed the presence of multiple variants of CyKILR as depicted in **Figure 2A,B**. To categorize these variants, we employed a classification based on the status of Exon 3. Those variants containing Exon 3 were designated as CyKILRa, while those lacking Exon 3 were denoted as CyKILRb. To evaluate their relative expression patterns, we conducted RT-PCR across a panel of 12 NSCLC lines. As shown in **Figure 2C**, expression of the CyKILRa group was easily detected in H358, H522, H2009, H1299, H1792, and H1648 cell lines. Moreover, a discernable level of expression was also observed in H2030, H1734, and H810 cell lines. In contrast, H460, A549, and H838 showed minimal to negligible expression of the CyKILRa group, the cell lines completely devoid of both CDKN2A and STK11 gene products as assessed by Western immunoblotting (**Fig.1G**). Conversely, the cohort of CyKILRb displayed expression exclusively within the H358, H522, H2009, H1299, and H1792 cell lines (**Fig.2C**), NSCLC cell lines with discernable expression of both CDKN2A and STK11 gene products by Western immunoblotting (**Fig.1G**). To quantify the expression level of CyKILRa and CyKILRb across the 12 NSCLC lines, we employed customized quantitative primers and probes (**Fig.2D**). To target Exon 3 inclusion variants (CyKILRa), the primers, and probes were designed to be complementary to Exon 3 (**Fig.2D**). In contrast, for the identification of Exon-exclusive variants (CyKILRb), the forward primer corresponded to Exon 1, while the reverse primer aligned with Exon 5, complemented by a probe spanning the junction of Exon 1 and Exon 5 (**Fig.2D**). Our results revealed that both CyKILRa and CyKILRb were predominantly expressed in H358, H522, H2009, H1299, and H1792 cell lines (**Fig.2D**), which are characterized by the presence of both active and WT *CDKN2A* and *STK11* genes (Expression of gene products derived from WT *CDKN2A* and WT *STK11* NSCLC lines verified by Western immunoblotting (**Fig.1G**)) vs other NSCLC cell lines, p = 0.000729 for n=4/cell line using One Way ANOVA). These data also demonstrate that CyKILRb has more distinct expression linked to these tumor suppressing genes.

**Figure 2.**
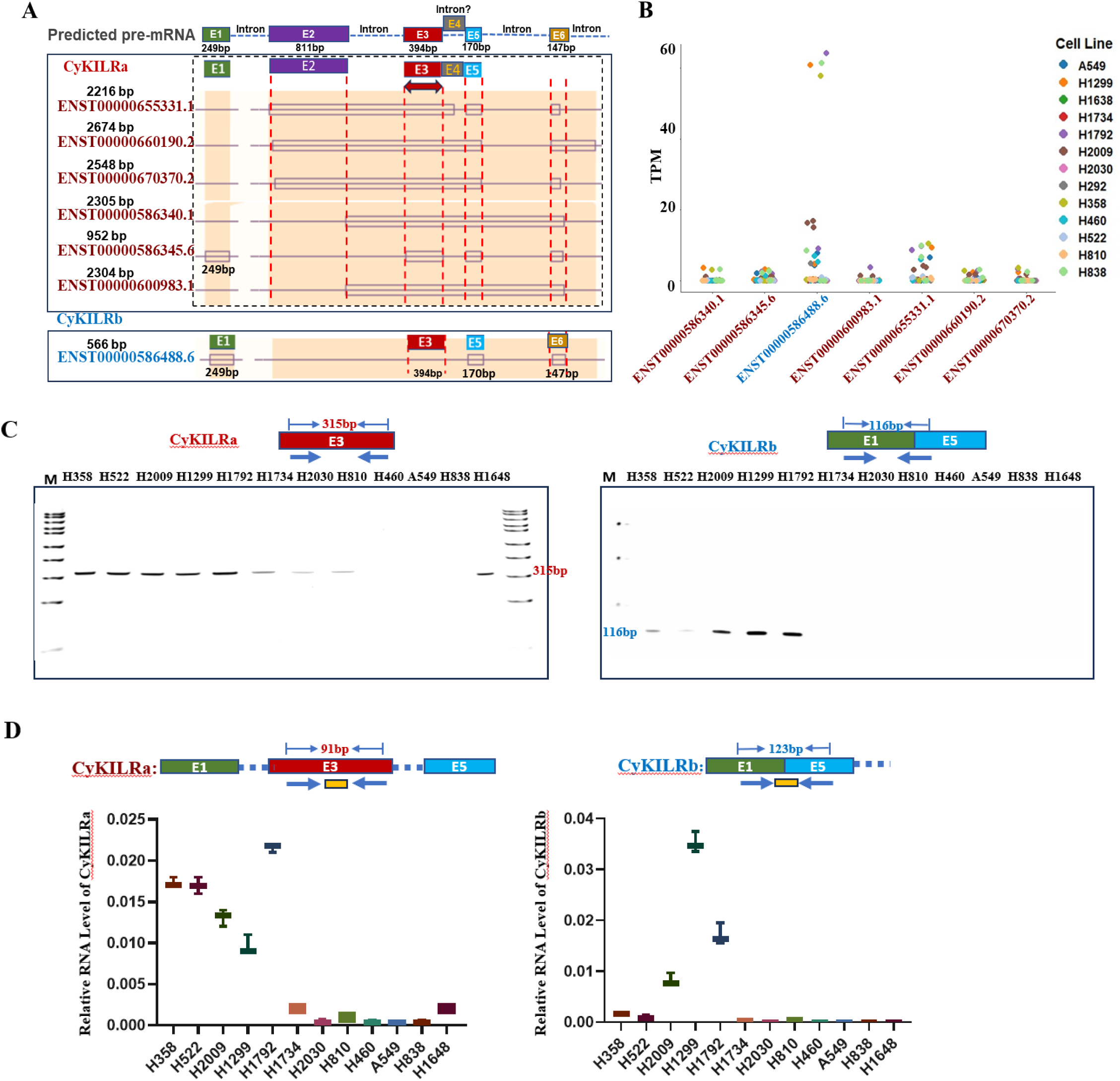
CyKILR is characterized by two distinct isoform groups based on exon 3 inclusion/exclusion. Total RNA was collected from the indicated NSCLC cell lines. Short-read deep RNA sequencing was conducted on these RNA samples. The R program was used for analysis of deep RNA sequencing data. **A.** The panel depicts the predicted pre-mRNA structure of CyKILR based on data from Ensembl. The CyKILR RNA splice variants were categorized into two distinct groups based on the inclusion/exclusion of Exon 3 into the mature RNA transcript. Predicted Exon and full-length CyKILR splice variants are annotated. Of note, the designated exon 6 is predicted to have multiple 5’ splice sites for several CyKILRa variants. **B.** The expression level of CyKILR isoforms across the 12 cell lines from the deep RNA sequencing conducted on RNA samples from these cell lines. The R program was used for analysis of deep RNA sequencing data. **C.** Relative RNA levels of CyKILRa (including Exon 3) and CyKILRb (excluding Exon 3) variants across a panel of 12 NSCLC cell lines using RT-PCR. **D.** The RNA expression levels of CyKILRa and CyKILRb are depicted by employing a customized quantitative primer and probe strategy for the detection of CyKILRa and CyKILRb variants by qRT-PCR and normalized to GAPDH mRNA levels. The data in Panel D are presented as mean□(SD) (n□≥□3 replicates). -RT controls were included in all experiments and replicates with no signal observed.

### CyKILRa and CyKILRb RNA splicing variants display differential cellular localization

Unlike mRNA, lncRNA is frequently detected in both the cytoplasm and nucleus. Research in the area suggests that an “AGCCC” motif plays a crucial role in the nuclear localization of numerous lncRNAs.[31] Following a sequence analysis, the specific presence of a “AGCCC” motif in Exon 3 of CyKILR was identified (**Fig.3A**). This discovery prompted us to investigate the localization patterns of the two distinct CyKILR groups by isolating total RNA from both the nucleus and cytoplasm. Subsequent multiplex and quantitative RT-PCR analyses highlighted a prevalent nuclear distribution for the variants containing Exon 3 (CyKILRa) in stark contrast a predominant cytoplasmic localization observed for the variants lacking Exon 3 (CyKILRb) (**Fig.3B,C**). Indeed, quantitative RT-PCR analyses showed >80% of CyKILRa variants clearly situated within the nucleus, while a corresponding percentage of >80% of CyKILRb variants consistently resided in the cytoplasm (**Fig.3C**). To validate our fractionation methodology, lncRNAs with reported cellular localizations were examined. CyKILRa localization correlated with the known nuclear-localized lncRNA, NEAT1, while CyKILRb localization correlated with the known cytoplasmic-localized lncRNA, TUG1 (**Fig.3D**).[32, 33] To determine whether the “AGCCC” motif of CyKILR exon 3 was regulating the nuclear localization, this sequence was removed by deletion mutagenesis. Ectopic expression of WT CyKILRa in NSCLC cells showed a high nuclear CyKILRa/cytoplasmic CyKILRa ratio showing the expected predominant nuclear localization (**Fig.3E**). In contrast, ectopic expression of a CyKILRa mutant lacking the “AGCCC” motif (Del-Mut) demonstrated a dramatic and significant reduction in the nuclear CyKILRa/cytoplasmic CyKILRa ratio (**Fig.3E**). These data demonstrate that the presence or absence of exon 3 in the mature CyKILR transcript regulates the cellular localization of this lncRNA via the “AGCCC” motif.

**Figure 3.**
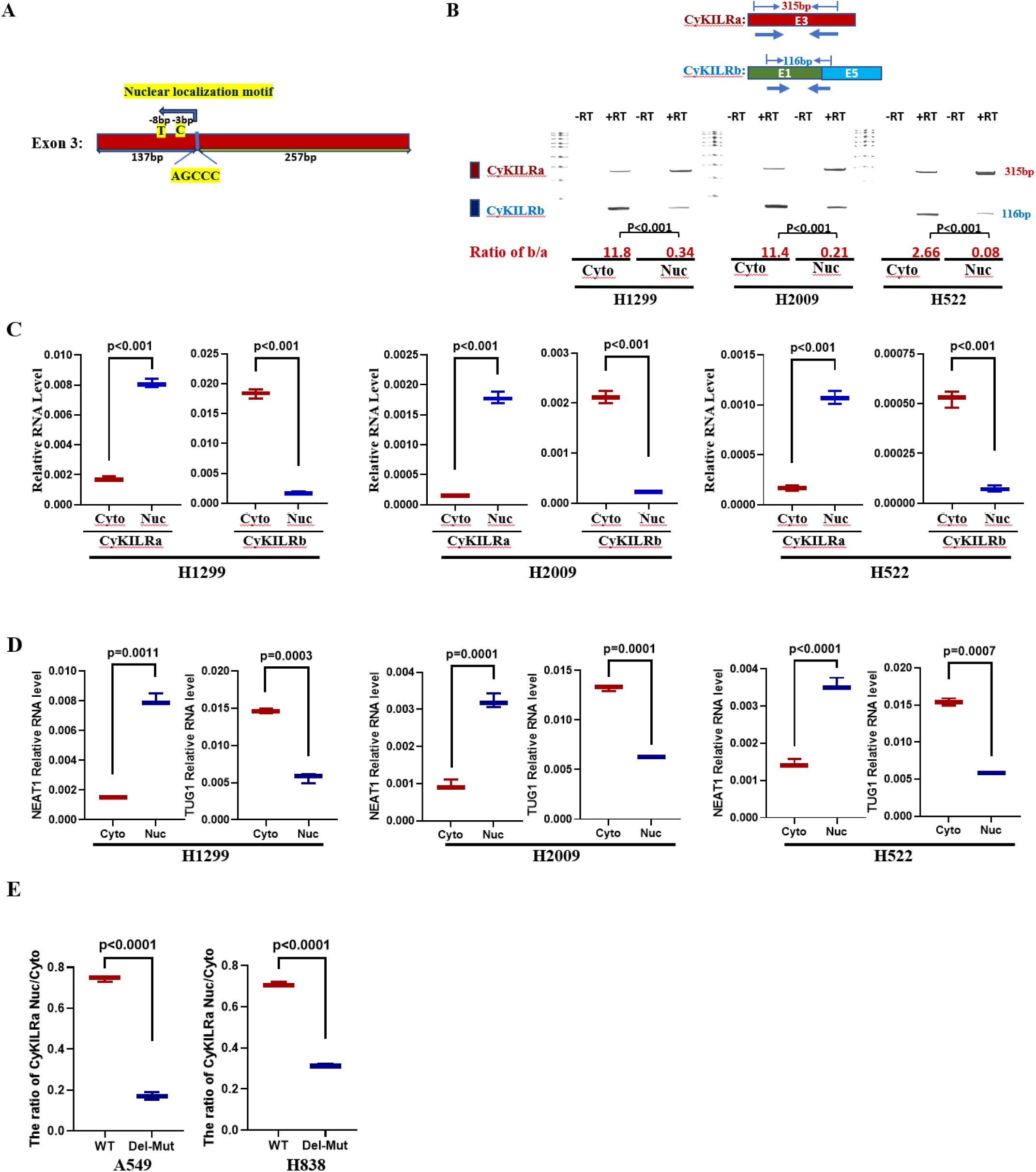
CyKILRa and CyKILRb demonstrate differential cellular localization regulated by an Exon 3 “AGCCC” motif. **A.** Sequence analysis of exon 3 of CyKILR reveals a nuclear localization motif, “AGCCC”. **B-D.** NSCLC cells from the depicted cell lines were fractionated for nuclei and cytoplasm. Total RNA was extracted from these fractions, reverse transcribed (RT), and subjected to multiplex PCR (**B**), quantitative PCR (qPCR) analysis for CYKILRa and b (**C**), or qPCR for NEAT1 and TUG1 (**D**). The data are presented as boxplots (n□= 3 replicates). **E.** Wild-type (WT) or a “AGCCC” deletion mutant of CyKILRa (Del-Mut) were ectopically expressed in two non-CyKILR-expressing cell lines (A549 and H833). Cells were fractionated for nuclei and cytoplasm. Total RNA was extracted from these fractions, reverse transcribed (RT), and subjected to qPCR) analysis for CYKILRa. The ratio of nuclear CyKILRa/cytoplasmic CyKILRa for WT and Del-Mut are depicted. Data in Panels B-E are presented as boxplots (n□= 3 replicates) and were analyzed by unpaired t-test with Welch’s correction. -RT controls were included in all experiments and replicates with no signal observed.

### CDKN2A and LKB1 regulate CyKILR exon 3 exclusion

Our data showing a strong correlation between WT, active *CDKN2A* and *STK11* genes with CyKILR expression suggested that the gene products of these two tumor suppressors cooperated to “drive” CyKILR expression. Surprisingly, downregulation of mRNAs stemming from the CDKN2A and STK11 genes, individually or concomitantly, by siRNA (**Fig.4A**) did not yield consistent or conclusive effects on the total CyKILR RNA levels in NSCLC lines although significant reduction in CyKILR expression was observed in non-cancerous, primary human bronchial epithelial cells (HBECs) (**Fig.4A,B**). In additional mechanistic studies, the RNA levels of the two splice variant groups of CyKILR (CyKILRa and CyKILRb) were examined individually, and whereas, individual downregulation of either CDKN2A or STK11 mRNAs by siRNA did not yield significant changes in the ratio between the two splice variant groups, the co-downregulation of the mRNA products of these two tumor suppressors led to a dramatic increase in the ratio of CyKILRb/CyKILRa variants compared to control siRNA (**Fig.4C**). These data demonstrate that LKB1 and CDKN2A gene products act to induce the inclusion of exon 3 into CyKILR RNA, which will enhance the expression of nuclear CyKILRa.

**Figure 4.**
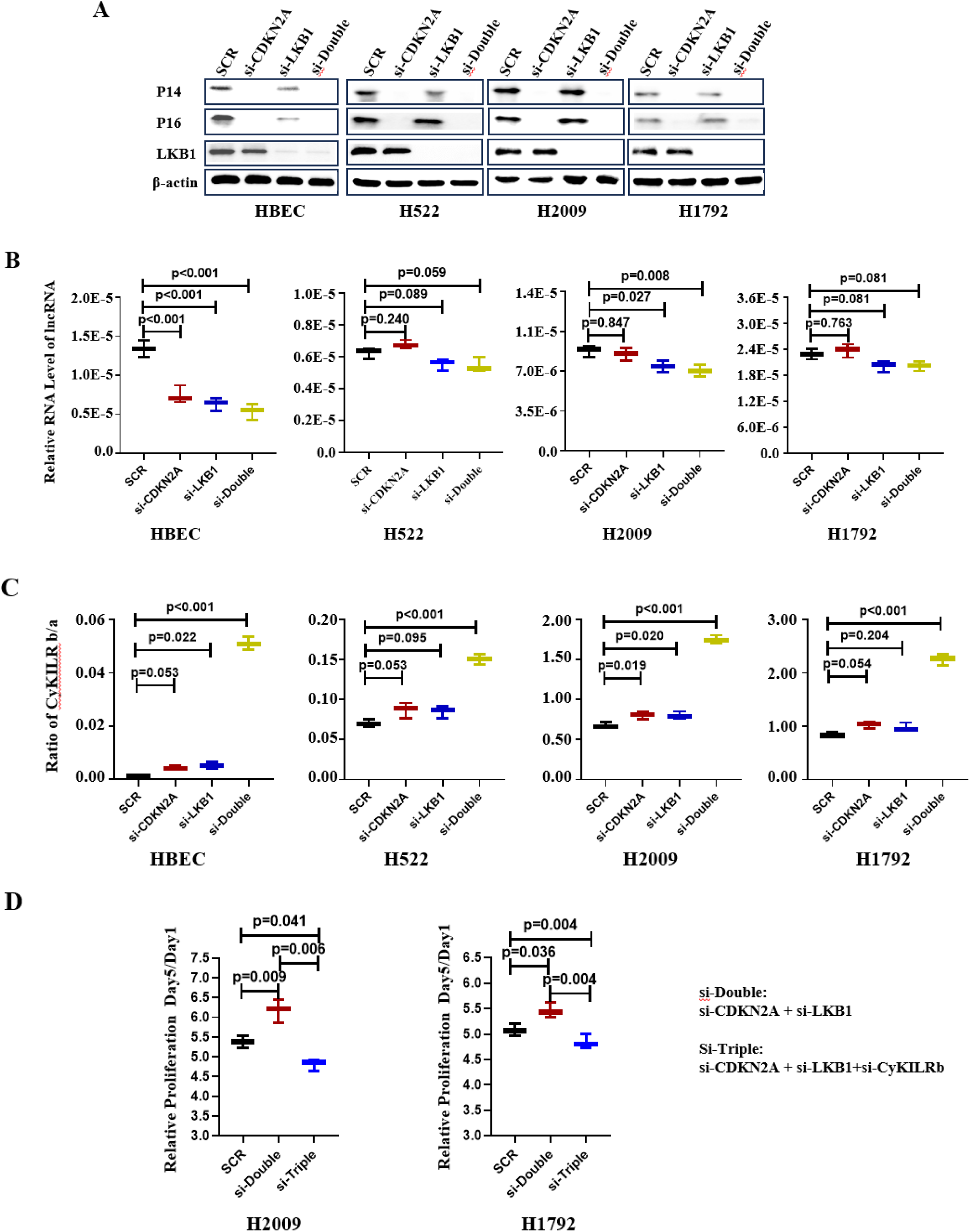
CDKN2A and LKB1 concomitantly regulate the inclusion of exon 3 into CyKILR RNA. The depicted cell lines were transfected with scrambled control (SCR), CDKN2A, LKB1, or CDKN2A/LKB siRNAs. After 48hrs, total protein and RNA were extracted. **A.** Protein levels of CDKN2A (p14 and p16) and LKB1 in samples subjected to single or double gene siRNA downregulation compared to scrambled siRNA controls (SCR). **B.** The level of total CyKILR RNA was determined by qRT-PCR normalized to GAPDH in samples subjected to single or double gene siRNA downregulation compared to scrambled siRNA controls. **C.** The ratio of CyKILRb to CyKILRa RNA levels determined by qRT-PCR in samples subjected to single or double gene siRNA downregulation compared to scrambled siRNA controls. **D.** The fold change of proliferation analysis of CyKILRb silencing on double gene downregulation by siRNA (CDKN2A and LKB1) compared to scrambled siRNA controls (SCR) post-48hr silencing (Day 0). Data in Panels B-D were analyzed by one-way ANOVA with Tukey’s multiple comparisons test and presented as box plots (n□≥□3 replicates). Data in Panel A are representative of n=3. -RT controls were included in all experiments and replicates with no signal observed.

The above findings inferred that induction of exon 3 inclusion into the mature CyKILR RNA transcript was biologically pertinent in relation to functional *CDKN2A* and *STK1*1 genes. To investigate whether suppression of CyKILRb expression affected biological phenotypes suppressed by the CDKN2A and STK11 gene products, proliferation was assessed when these tumor suppressors were again co-downregulated by siRNA. Downregulation of both CDKN2A and STK11-derived mRNAs induced a significant increase in cell proliferation (**Fig.4D**). The simultaneous downregulation of CyKILRb attenuated the increased proliferation observed following CDKN2A and LKB1 co-silencing as compared to control siRNA cells (**Fig.4D**). These data show that suppression of CyKILRb expression via inclusion of exon 3 is an important mechanism by which LKB1 and CDKN2A subdue cell proliferation. To further interrogate these biological findings, all CyKILR transcripts were downregulated in multiple NSCLC cell lines with WT *STK11* & *CDKN2A* genes, which required the utilization of DNA/RNA chimeric antisense oligonucleotides (ASOs) targeting intron 1 of the pre-mRNA to silence nuclear CyKILR as siRNA has been shown ineffective for these nuclear forms of RNA[34]. Downregulation of CyKILR pre-mRNA induced significant increases in cellular proliferation (**Figure S2**), and thus, CyKILR, in general, is a suppressor of cell growth and proliferation, while the CyKILRb splice variant possesses an opposing function.

### Nuclear CyKILRa and cytoplasmic CyKILRb have opposing functions *in vitro* and *in vivo*

Our data suggested that CyKILR has biological functions linked to cancer cell phenotypes (e.g., cell proliferation), which may also be opposing depending on alternative RNA splicing and cellular localization. Thus, the role of the individual CyKILR isoform groups was examined for effects on cell proliferation, cell clonogenic survival, and cell migration. First, the functional role of nuclear-localized CyKILR variants (CyKILRa) was assessed, which again required the utilization of DNA/RNA chimeric antisense oligonucleotides (ASOs) to silence nuclear CyKILRa, as siRNA has been shown ineffective for these nuclear forms of RNA.[34] The silencing efficacy of our ASOs exceeded 50% across all four tested cell lines (**Fig.5A**). Downregulation of CyKILRa resulted in significant enhancement of cell proliferation (**Fig.5B**), and a >two-fold acceleration in cell migration (**Fig.5C**). Furthermore, cell clonogenic survival was significantly increased in these cell lines by downregulation of CyKILRa (**Fig.5D**). In addition, downregulation of CyKILRa had a significant and profound effect on subcutaneous (sc) tumor formation in mice for both the H2009 and H1792 cell lines. Specifically, downregulation of CyKILRa with ASOs induced significant tumor growth and formation for both cell lines injected sc in the flanks of mice after four weeks, a time point when tumors were not appreciably formed in the control cell lines (**Fig.5E**). These cellular effects could not be attributed to artifactual or regulatory effects on the expression of genes located in close proximity to the CyKILR transcription locus (**Figure S1**) as downregulation of CyKILRa had no effect in the expression of these genes (**Figure S3**). In stark contrast to CyKILRa, downregulation of CyKILRb by siRNA (**Fig.6A**) resulted in a significant reduction of cell proliferation (**Fig.6B**). Additionally, a significant decrease in both cell migration and cell clonogenic survival was observed (**Fig.6C,D**). Downregulation of CyKILRb by siRNA in H2009 and H1792 cells also induced a significant reduction in sc tumor growth in mice when control tumors were allowed to achieve at least 200 mm^3^ in volume (**Fig.6E**). In all replicates, downregulation of CyKLRa and b produced profound effects on tumor growth over the experimental time course (**Figure S4**). These data demonstrate that nuclear CyKILRa has specific cellular functions and acts to suppress key pro-survival phenotypes observed in NSCLC cells. Furthermore, cytosolic CyKILRb has an opposing function and acts as a pro-survival, pro-tumorigenic, and pro-growth factor.

**Figure 5.**
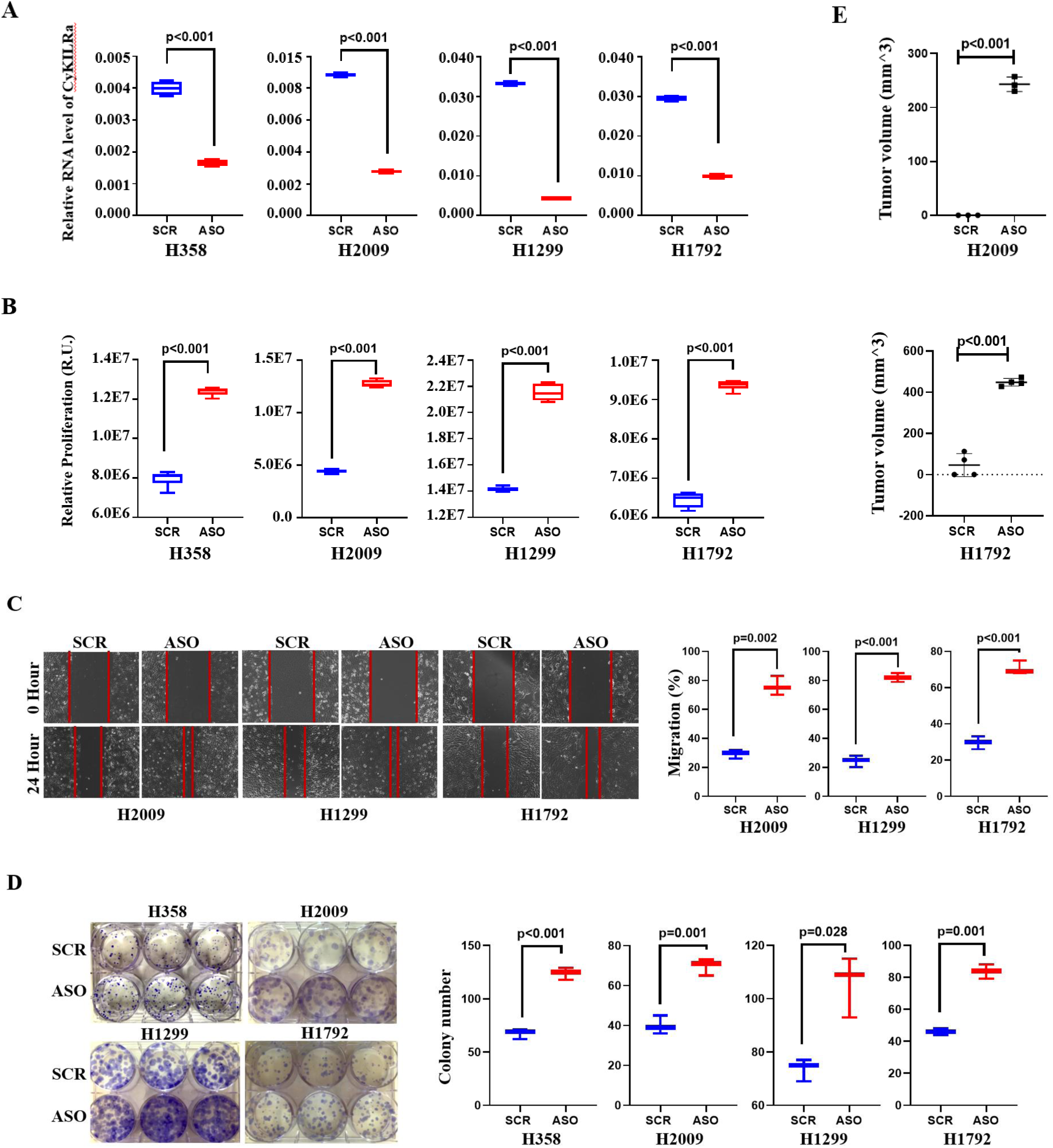
Nuclear CyKILRa acts as a tumor suppressing factor. The depicted cell lines were transfected with scrambled control ASO (SCR) or CyKILRa ASO (ASO). After 48hrs, total RNA was extracted or in concomitant experiments, the cells were assessed for the indicated biological mechanisms with the 48hr post-transfection acting as the 0 time point (**A-D**). In addition, 48hr post-transfected H2009 and H1792 cells were subsequently mixed with Matrigel in a 1:1 ratio and injected subcutaneously (sc) into the flanks of five-week-old female NOD-SCID mice (1 x 10^6^ cells per mouse per flank). The left flank received Control-treated cells, while the right flank received knockdown cells (**E**). **A.** RNA levels of CyKILRa in the depicted cell lines post-ASO treatment normalized to the mRNA levels of GAPDH. -RT controls were included in all experiments and replicates with no signal observed. **B.** Proliferation of CyKILRa ASO-treated cells compared to scrambled control ASO-treated cells at the 48-hour time point. **C.** Migration of CyKILRa ASO-treated samples compared to scrambled control ASO-treated cells at the 24-hour time point. **D.** Plate clonogenic survival of CyKILRa ASO-treated cells compared to scrambled control ASO-treated cells. **E.** Tumor volume of ASO-Con or CyKILRa ASO treated H2009 or H1792 in mice. The data are presented as box plots (n□≥□3 replicates), and statistical analysis was performed using the unpaired t-test with Welch’s correction.

**Figure 6.**
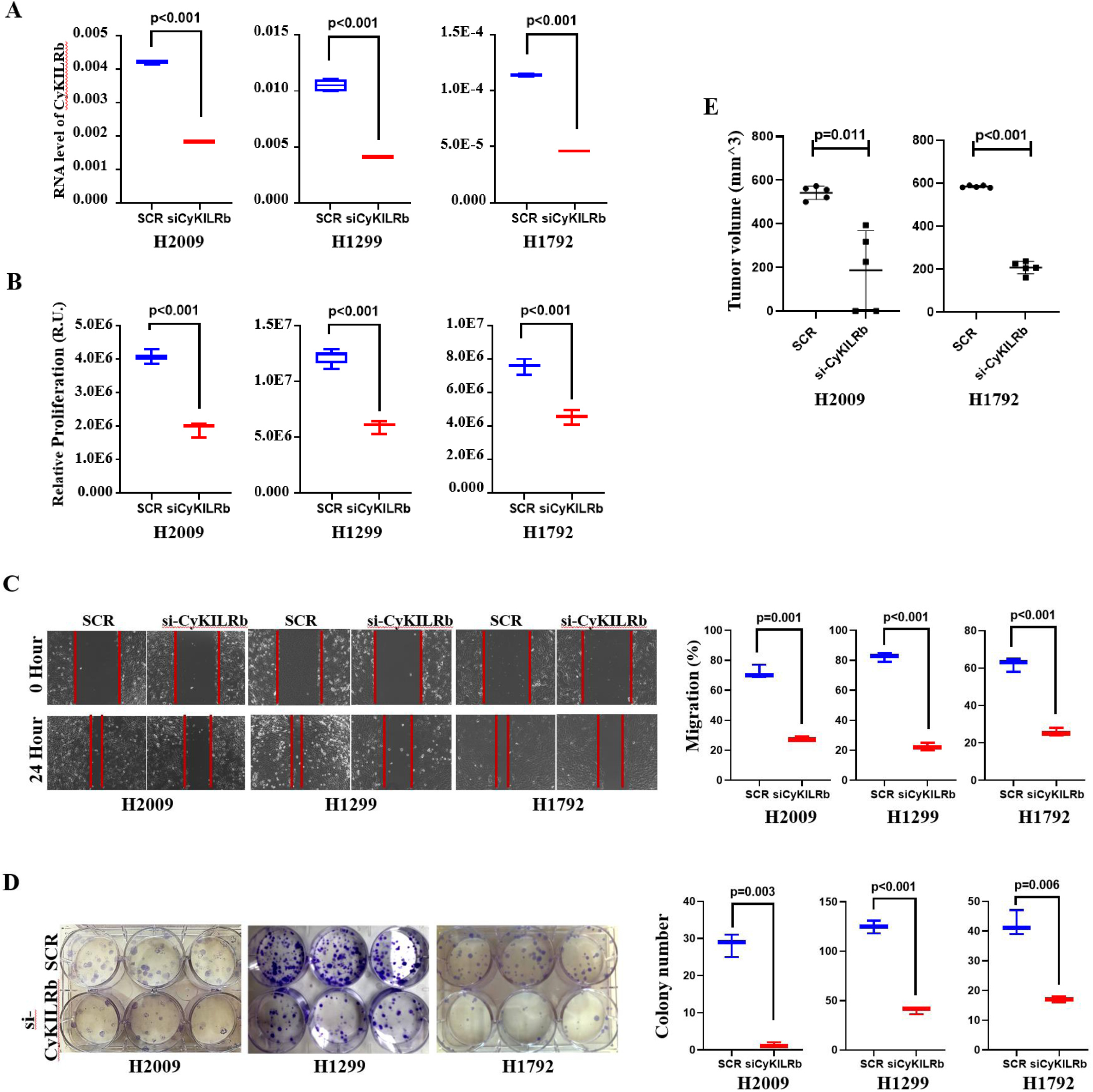
Cytoplasmic CyKILRb is a prosurvival factor. The depicted cell lines were transfected with scrambled control (SCR) or CyKILRb siRNA (siCyKILRb). After 48hrs, total RNA was extracted or the cells were assessed for the indicated biological mechanisms with the 48hr post-transfection acting as the 0 time point (**A-D**). In addition, 48hr post-transfected H2009 and H1792 cells were subsequently mixed with Matrigel in a 1:1 ratio and injected subcutaneously (sc) into the flanks of five-week-old female NOD-SCID mice (1 x 10^6^ cells per mouse per flank). The left flank received Scramble-treated cells, while the right flank received knockdown cells (**E**). **A.** RNA levels of CyKILRb siRNA-treated samples compared to scrambled siRNA control (SCR) samples normalized to GAPDH mRNA. -RT controls were included in all experiments and replicates with no signal observed. **B.** Proliferation of CyKILRb siRNA-treated cells compared to scrambled siRNA control cells at the 48-hour time point. **C.** Migration of CyKILRb siRNA-treated cells compared to scrambled siRNA control cells at the 24-hour time point. **D.** Plate clonogenic survival of CyKILRb siRNA-treated cells compared to scrambled siRNA control cells. **E.** Tumor volume of siRNA-scramble control or CyKILRb siRNA treated H2009 or H1792. The data are presented as box plots (n□□≥□□3 replicates), and statistical analysis was performed using the unpaired t-test with Welch’s correction.

### CyKILRa and CyKILRb have opposing functions on cell signaling pathways

Since modulation of CyKILR splice variants showed strong effects on tumor biology and cancer cell phenotypes, the mechanistic underpinnings were explored. In this regard, downregulation of CyKILRa and b followed by deep RNA sequencing and pathway analysis was undertaken. Downregulation of CyKILRa by ASOs showed decreased expression of factors central in apoptosis induction (e.g., Bak1, CHOP, PUMA) as well as significantly increased expression of factors known to drive the cell cycle, cell proliferation, and cell growth (e.g., CDK1, CDK2, CHEK2) (**Fig.7A-C; Figures S5,S6,S7A-D**). Downregulation of CyKILRb, on the other hand, led to induction of pro-survival pathways (**Figure S7E)** and tumor suppressing miRNAs, hsa-miR-424-3p and has-miR-503-p (**Fig.7D-F**). Treatment of H2009 and H1299 cells with mimics of these two miRNAs partially, but significantly suppressed key cancer cell proliferation (**Fig.7G**). Highlighting the complexity of CyKILR in the regulation of cell signaling and gene expression, CyKILRb also serves as a suppressor of CyKILRa expression as downregulation of CyKILRb led to a significant increase in CyKILRa expression (**Figure S8**). This is in stark contrast to CyKILRa as ASO downregulation of this isoform did not significantly affect CyKILRb expression (**Figure S8**). These data show that CyKILR splice variants have complex roles in the modulation of key pathways linked to both tumor suppression and tumorigenesis.

**Figure 7.**
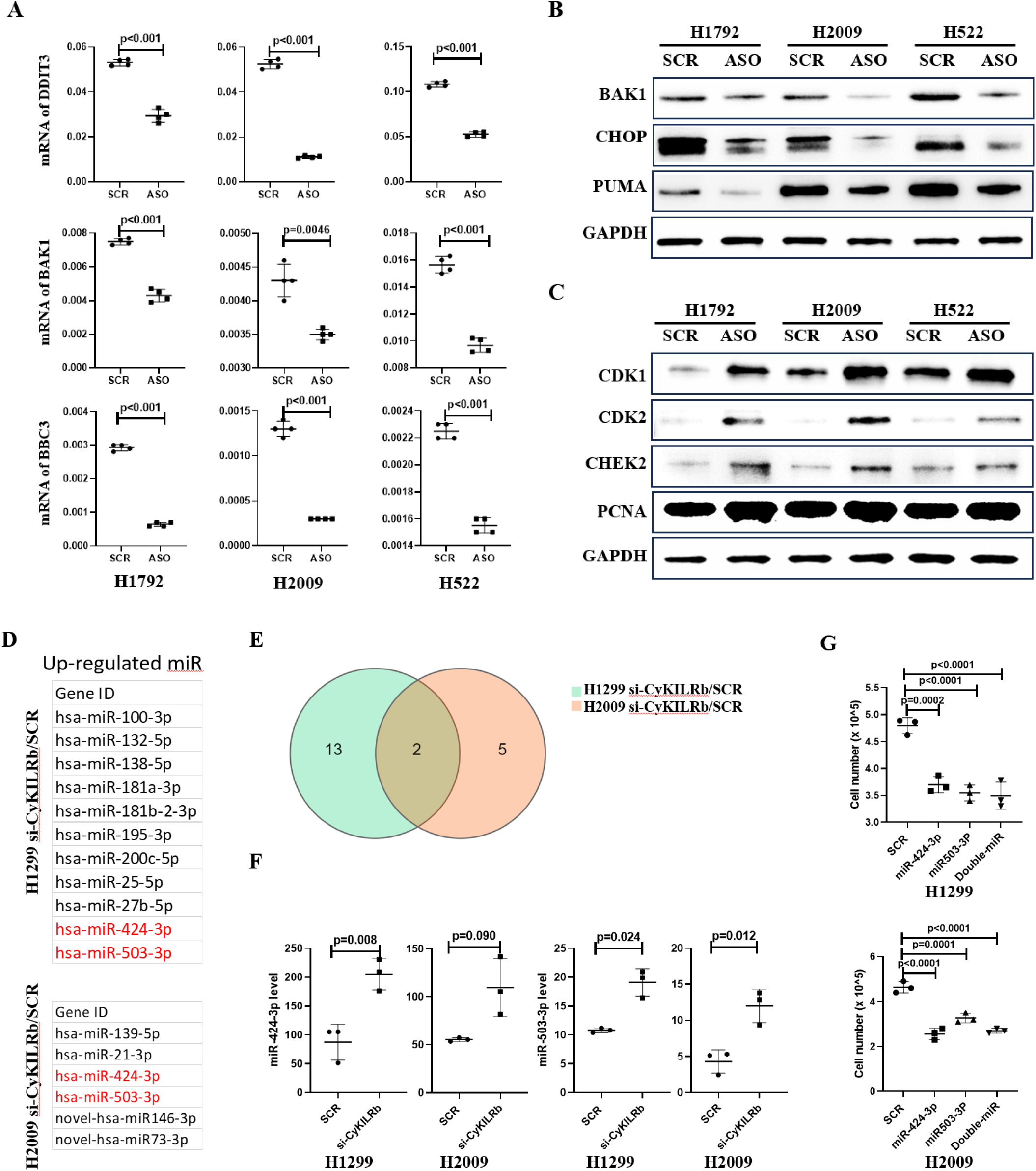
CyKILRa and CyKILRb apposingly regulate cell growth signaling pathways. H2009 or H1792 cells were treated with either scrambled control ASO (SCR) or CyKILRa ASO (ASO), and H1299 and H2009 cells were treated with either siRNA-Con (SCR) or CyKILRb siRNA (siCyKILRb). Total RNA from cell subjected to these treatments underwent deep RNA sequencing. Following data filtering and clean read alignment, gene expression levels and differential expression gene analysis were conducted using RSEM (v1.2.28) and DESeq2. KEGG enrichment analysis of annotated differentially expressed genes was performed using R Phyper. The CyKILRa ASO-treated groups exhibited regulation of cell apoptosis and cell cycle pathways compared to the control-treated groups (**A-C**). While the CyKILRb siRNA-treated groups exhibited the regulation of microRNAs (**D-G**). **A.** RNA levels of key apoptosis markers were down-regulated by CyKILRa ASO in all cell lines as assessed by qRT-PCR. -RT controls were included in all experiments and replicates with no signal observed. **B.** Protein levels of key apoptosis markers were down-regulated by CyKILRa ASO in all cell lines as assessed by Western immunoblotting. **C.** Protein levels of key cell cycle markers were up-regulated by CyKILRa ASO in all cell lines. **D.** Up-regulated microRNAs by CyKILRb knockdown in both cell lines. **E.** The Venn diagram displays the upregulated miRNAs resulting from CyKILRb knockdown in both H1299 and H2009 cell lines. **F.** RNA levels of miR-424-3P and miR-503-3P in H1299 and H2009 cells treated with si-CyKILRb vs scramble control (SCR). **G.** Cell proliferation assay comparing miR-424-3P and miR-503-3P mimic-treated H1299 and H2009 cells vs scramble-treated (SCR) cells 48 hours post-transfection. The data in Panels A, F, and G are presented as box plots (n□□≥□□3 replicates), and statistical analysis was performed using the unpaired t-test with Welch’s correction. Data in Panels B and C are representative of n = 3.

## DISCUSSION

In this study, we have discovered a novel long non-coding RNA (lncRNA), which we termed **<u>Cy</u>**clin-Dependent **<u>K</u>**inase **<u>I</u>**nhibitor 2A-regulated **l**nc**<u>R</u>**NA (CyKILR) due to its expression being mainly confined to NSCLC lines harboring a wild-type and active *CDKN2A* gene. Additional studies showed that an oncogenotype consisting of both a WT *CDKN2A* and *STK11* gene correlated stronger with the expression of CyKILR, and downregulation of the mRNA of either gene led to significant loss of CyKILR expression in primary HBECs. The loss of CyKILR expression observed in NSCLC cell lines lacking either a WT STK11 or CDKN2A gene correlated and corroborated this finding, which strongly suggested that CyKILR has a possible tumor suppressor role linked to the cell signaling pathways activated by the LKB1 and CDKN2A gene products. In support of this conclusion, downregulation of the CyKILRa isoforms (exon 3 included) significantly enhanced cell clonogenic survival, migration, and proliferation as well as tumor growth. On the other hand, downregulation of CDKN2A and STK11 mRNAs in NSCLC cell lines with both WT STK11 and CDKN2A genes surprisingly demonstrated minimal or no significant effect on total CyKILR expression. Thus, depending on the cellular context and oncogenic landscape, maintenance of CyKILR expression becomes independent of these genes. Shedding light on a plausible adaptive mechanism was the additional characterization of CyKILR, which identified multiple RNA splice variants including an exon 3 excluded isoform designated CyKILRb. In contrast to CyKILRa, this splice variant class was pro-survival, and notably, the ratio of CyKILRb/a, was very low in non-cancerous, HBECs, when compared to the NSCLC cell lines that expressed CyKILR. We posit that shifting from the CyKILRa isoform to the CyKILRb isoform through modulation of exon 3 inclusion/exclusion by alternative RNA splicing is required for transformed NSCLC cells to maintain cell homeostasis in the presence of these key tumor suppressors. Indeed, downregulation of CyKILRb had dramatic effects in reducing the clonogenic survival and tumor growth of NSCLC cells with WT STK11 and CDKN2A genes. Specific downregulation of CyKILRb also blocked any enhancement of cellular proliferation observed when both of these genes were inactivated. Therefore, the CyKILRb splice variants of this lncRNA are plausible proto-oncogenes, or more specifically, these data suggest that exclusion of exon 3 from CyKILR RNA removes a key nuclear lncRNA required for LKB1 and CDKN2A gene products to act as efficient tumor suppressors.

The observation that the STK11 and CDKN2A gene products co-regulated the alternative RNA splicing of CyKILR was a unexpected finding. Indeed, a tremendous 30-fold increase in the CyKILRb/a RNA ratio in HBECs was observed when CDKN2A and STK11 mRNAs are downregulated. This is a 10- fold higher response than the NSCLC cells examined in this study. These data suggest that transformed cells with an active and WT CDKN2A and STK11 gene also have active oncogenic pathways with opposing function on downstream RNA splicing, which act to suppress exon 3 inclusion. How the products of these tumor suppressor genes regulate the inclusion/exclusion of CyKILR exon 3 is unknown opening avenues of unanswered questions in relation to their tumor suppressing functions and the regulation of alternative RNA splicing. Dysregulated alternative RNA splicing is well established in cancer with RNA *trans*-factors acting as proto-oncogenes,[35] but there is essentially nothing reported in the literature as to alternative RNA splicing regulation downstream of the signaling pathways regulated by CDKN2A and STK11 gene products. There are reported links to the CDKN2A gene product, p16, in regulating the association of the RNA binding protein, AUF1 to CDKN1A mRNA in regard to RNA stability.[36] The expression of hnRNP B1, an RNA splicing factor, was reported as inversely regulated in relation to p16 expression in lung cancer tissue.[37]. CDKN2A gene products as well as LKB1 will repress activation of the AKT pathway, and thus, when CDKN2A and LKB1 are downregulated, AKT activation will occur, which is known to have distinct effects on RNA splicing.[38–41] This signaling paradigm is logical as loss of both pathways may cooperate to enhance the expression of specific AKT isoforms, modulate specific post-translational modifications related to AKT activation, and alter cellular topology via PIP_3_ production in specific organelles such as the nuclear membrane or nucleus itself. As to the latter, it has been previously surmised and shown that AKT can directly phosphorylate RNA *trans*-factors in the nucleus to modulate specific RNA splicing events.[38–40] Thus, coordinated modulation/suppression of nuclear AKT signaling may be one plausible route for loss of these tumor suppressors to cooperate in regulating specific alternative splicing events. Additionally, our study has not examined the alternative RNA splicing in the NSCLC cells lacking appreciable CyKILR expression. Thus, downregulation and suppression of total CyKILR expression may occur via the induction of CyKILR RNA splice variants removed by nonsense mediate RNA decay[42], which is an efficient mechanism to remove aberrant mRNA transcripts that may arise from alternative RNA splicing. Future studies are required to determine how loss of the *STK11* and *CDKN2A* tumor suppressors leads to suppression of CyKILR expression in NSCLC.

The characterization on CyKILR RNA splice variants in this study led to the observation of distinct subcellular distributions for CyKILRa and b, which corresponded to their divergent functions. Indeed, lncRNAs serve as operational units/scaffolds, and their subcellular localization plays a pivotal role in governing their functions.[43] For example, a substantial number of lncRNAs primarily reside within the nucleus, where they perform diverse regulatory functions through a multitude of combinatorial mechanisms optimized for nuclear enrichment.[44–47] In our study, we found that CyKILRa has a distinct nuclear localization in contrast to the cytoplasmic predominant localization of CyKILRb. Through in silico analysis, a potential nuclear retention motif, “AGCCC,” was identified with -8 (T or A) and -3 (G or C) relative to the first nucleotide of the pentamer in Exon 3 of CyKILR[31]. Thus, CyKILRa predominance in localizing to the nucleus in contrast to CyKILRb is logical as this sequence only resides in exon 3, and our empirical experimental results validated the requirement of this motif in regulating CyKILRa nuclear localization. As to how CyKILRa functions as a tumor suppressor, its nuclear localization may be key to exerting regulatory influence over cellular processes, possibly through interactions with epigenetic factors involved in histone modification complex scaffolding or recruitment as previously reported for other lncRNAs.[48] Additionally, nuclear lncRNAs often assume additional roles including the modulation of pre-mRNA processing and involvement in alternative splicing events.[49] Uncovering the specific regulatory mechanisms governing CyKILRa within the nucleus holds great promise for precision-targeted tumor therapy as our transcriptomics analyses show that AKT levels were significantly increased in two separate NSCLC cells lines when CyKILRa was downregulated by ASO, and this pathway is potent in both survival signaling, oncogenesis, and in modulating alternative RNA splicing. Indeed, systems level analyses of the transciptome showed the apoptosis genes were downregulated and other pathways linked to enhancing the cell cycle were increased, which our laboratory confirmed by qRT-PCR and Western immunoblotting. These data support the inference that CyKILRa acts as a nuclear scaffold for regulating gene expression, specifically suppression of cell cycle pathways while simultaneously maintaining the levels of factors allowing for apoptosis to occur upon oncogenic stress.

In contrast to CyKILRa, CyKILRb functions as a pro-tumorigenic factor, and like CyKILRa, it is currently unknown as to how CyKILRb functions in this role. Emerging research also suggests that the population of cytoplasmic lncRNAs may be more abundant than previously recognized.[50–52] Cytoplasmic lncRNAs have been implicated as a microRNA sponge, sequestering microRNAs or regulating protein stability.[53] When lncRNAs function as miRNA “sponges”, these non-coding RNAs act as competing endogenous RNAs (ceRNAs),[54] which are reported to play key roles in cancer disease states.[55] In this regard, systems level analysis of CyKILRb downregulation in two NSCLC cell lines with WT *STK11* and *CDKN2A* genes found a consistent increase in two reported, tumor suppressing miRNAs, hsa-miR-424-3p and has-miR-503-p.[56, 57] Interestingly, there is no complementary sequence for these miRNAs in CyKILRb suggesting that the regulation of these miRNAs by CyKILRb is not through a “sponge” mechanism, but possibly via modulation of cell signaling pathways affecting the expression of these miRNAs. Indeed, CyKILRb may act as a “sponge” for other miRNAs that then regulate cell signaling pathways modulating the expression of these tumor suppressing miRNAs analogous to other cytoplasmic lncRNAs like TUG1.[33] Regardless, examining this novel link between CyKILRb and the expression of hsa-miR-424-3p and has-miR-503-p is a future mechanistic avenue to explore.

In conclusion, our study characterizes a novel lncRNA with functional roles in NSCLC with a specific oncogenotype. The association with CDKN2A and LKB1 status, the differential subcellular localization, and the impact on cellular behavior and in vivo tumorigenicity suggest that CyKILR splice variants are important players in NSCLC pathogenesis. Further research is needed to elucidate the precise molecular mechanisms underlying the functions of the individual CyKILR variants and their expression. Indeed, this intricate interplay between the two subcellular groups of CyKILR underscores the critical importance of their relative ratios offering a precise means to finely tune cellular progression. Additionally, exploration as to its potential as a therapeutic target in NSCLC and potentially other cancer types is warranted as RNA-based therapies are now plausible.[58] Overall, this study provides a foundation for future investigations into the diagnostic and therapeutic implications of targeting specific lncRNA splicing variants in NSCLCs, and future exploration of lncRNAs like CyKILR offers a promising avenue for precision medicine approaches in cancer therapy.

## MATERIALS AND METHODS

### Cell lines and materials

NCI-358, NCI-H522, NCI-H2009, NCI-H810, NCI-H460, A549, NCI-H838, NCI-H1734, NCI-H1648, NCI-H2030, NCI-H1299, NCI-H1792, and Primary human bronchial epithelial cells (HBECs) were purchased from ATCC. RNA Subcellular Isolation Kit was from Active Motif (25501). SuperScript™ IV VILO™ Master Mix with ezDNase™ Enzyme (11766050), TaqMan™ Universal PCR Master Mix (4324018), Opti-MEM™ I Reduced Serum Medium (31985062), Lipofectamine™ RNAiMAX Transfection Reagent (13778150), and Pierce Fast Western Blot Kit (35050) were from Thermo Fisher Scientific. Cell Titer□Glo 2.0 Assay Kit was purchased from Promega (G9242). The mycoplasma contamination detection kit was purchased from InvivoGen (rep-mys-50).

### Cell culture

All cells were cultured following the procedures described previously and were utilized within 6 months or -twenty passages, whichever came first, from the time of receipt and thawing. [59] Throughout the entire study, cell lines were subjected to Mycoplasma testing every two months using a Mycoplasma contamination detection kit. Parental cell lines were authenticated by STR (short tandem repeat) profiling.

### Transcriptomics and Splicomic analyses

*Short-Read:* RNA sequencing was conducted using 150 bp paired-end reads, aiming for a depth of approximately 90 million reads for each RNA sample. Sequencing reads were aligned to the GRCh38.p14 reference genome using STAR software (v2.7.10b), and post-alignment, BAM files were filtered and indexed with Samtools (v1.16.1) to remove low-quality reads.[60, 61] For differential gene expression analysis, the R package DESeq2 was utilized on previously described RNAseq data processed with Salmon 1.10.0.[62, 63] Additional R packages employed included tximport for importing data, biomaRt for genomic annotation, tidyverse for statistical analyses, RColorBrewer for color palettes, and org.Hs.eg.db for gene annotations.[64–66] Isoform detection and quantification was performed using the previously described Salmon processed RNAseq data. Transcript IDs for genes of interest were identified from the GENCODE v45 comprehensive annotation .gtf file. These IDs facilitated parsing of Salmon .sf files to obtain TPM values for each transcript, which informed our transcript abundance calculations. Finally, TPM values for selected transcripts were visualized using the tidyverse package in R version 4.3.3.[64] Differential alternative splicing analysis was performed on these filtered BAM files using rMATS turbo (v4.1.2).[67] Subsequently, R programming language was utilized to filter and generate a list of splicing targets from the data produced by rMATS.

### RNA extraction and Quantitative RT-PCR (qRT-PCR)

Total RNA and subcellular RNA were isolated from cultured cells using the RNA Subcellular Isolation Kit from Active Motif (25501) following the manufacturer’s instructions with DNase treatment to remove genomic DNA contamination. RNA quantification was performed using a NanoDrop NP-1000 spectrometer, and RNA (2 μg) was used for the first-strand cDNA synthesis using SuperScript™ IV VILO™ Master Mix with ezDNase™ Enzyme. Subsequently, qRT-PCR was conducted as done on a Quant Studio 5 instrument (Applied Biosystems, Carlsbad, CA, USA). [38–40, 48, 49, 53, 68] Superscript “minus” (-RT) controls were included in all experiments to confirm a lack of target amplification caused by genomic DNA contamination. The housekeeping gene GAPDH was utilized for normalization using the ΔΔCt method. The specific primers and probes used in the study are listed in **Table S2**. A customized quantitative primer and probe strategy for the detection of CyKILRa and CyKILRb variants was also employed. For CyKILRa Exon 3 inclusion variants, we designed primers and probes specifically targeting Exon 3, as all CyKILRa variants in this group contain Exon 3. In contrast, for CyKILRb Exon 3 exclusion variants, which lack Exons 2-4, the forward primer aligns with Exon 1, the reverse primer is complementary to Exon 5, and a probe spans the Exon 1 to Exon 5 junction.

### Multiplex RT-PCR

A reaction mixture was prepared by combining cDNA (25 ng) with both sets of primers at a final concentration of 0.1 μM. To achieve a total volume of 50 μl, the cDNA-primer mixture was mixed with 2X master mix and water. The PCR was performed using the hot-start method, with an annealing temperature of 58°C for 26 cycles. The specific primers used in the study are listed in **Table S2**.

### siRNA and ASOs Transfection

Cells were plated in a full medium without antibiotics the day before transfection, ensuring they reached 50% confluency at the time of transfection. To dilute siRNA or ASOs, Opti-MEM® I Reduced Serum Medium was used. For siRNA transfection, Lipofectamine RNAiMAX was employed as done,[54, 55, 69–71] while Lipofectamine 3000 was used for ASOs transfection. The final concentration of both siRNA and ASOs in the cell culture medium was 20nM. The sequences of siRNAs and ASOs are listed in **Table S2**. ASO was customized 2′-*O*-methyl locked DNA chimera with phosphorothioate bonds backbone. siRNA was customized 19-22 basepair duplex RNA with 3’ UU overhangs.

### Plasmid of wildtype or deletion mutant of CyKILRa Transfection

The full-length CyKILRa (ENST00000586345.6) and the AGCCC deletion mutant were synthesized and inserted into the plasmid pcDNA3.1 between the Not I and Xho I sites. A total of A549 or H838 cells (1×10^6^) were seeded in a 10 cm plate and allowed to incubate overnight prior to transfection. Plasmids (5 µg) were transfected using Lipofectamine 3000 (12.5 µL) according to the manufacturer’s instructions. Subcellular RNA extraction was performed 48 hours after transfection.

### Proliferation analysis

To investigate cell proliferation, the Cell Titer-Glo Luminescent Cell Viability kit was utilized following the manufacturer’s instructions. Cells were initially seeded in 96-well plates at a density of 2000 cells per well in RPMI1640 supplemented with 10% FBS. After treatment with ASO or siRNA for 48 hours, the plates were carefully decanted and refilled with 50 μl of Cell Titer-Glo reagent and 50 μl of PBS per well. The plates were then incubated for 10 minutes at room temperature in the dark. Subsequently, the luminescence readings were taken using the Gene5 Multi-Mode Microplate Reader (Molecular Devices, Sunnyvale, CA) with a 250 ms integration.

### Cell clonogenic survival assays

After ASO or siRNA treatment, cells were harvested and re-suspended in complete growth media. Subsequently, the cells (2-4 x10^2^) were seeded onto six-well plates and allowed to grow until visible colonies formed as done,[38–40, 48, 68, 72–74], which typically occurred within 7-21 days depending on the cell line. Once the colonies became visible, they were fixed with cold methanol, stained with 0.25% crystal violet in 25% methanol, washed, and then left to air-dry. Any colony with greater than 40 cells/colony were counted.

### Cell Migration

The scratch migration assay was conducted following the procedure previously described.[75–78] Briefly, ASOs- or siRNA-treated cells were cultured in 6-well plates with RPMI 1640 medium containing 10% FBS until they reached 90% confluency. An ∼1 mm cell-free gap was created using a sterile pipette tip. After washing away dislodged cells with PBS, the plates were refilled with fresh medium and incubated for 24 hours before capturing images. Cell migration was quantified using Image J software, analyzing the change in the cell-free area at 0 hours and 24 hours.

### Western immunoblotting

Cells were lysed in RIPA buffer containing Halt™ Protease and Phosphatase Inhibitor Cocktail (Thermo Fisher Scientific, 78440). For protein concentration determination, the Pierce BCA Protein Assay kit (Thermo Fisher Scientific, 23227) was used.[49, 79–81] Whole-cell lysates were mixed with Laemmli loading buffer, boiled, separated by 4-20% SDS–PAGE, and transferred to a PVDF membrane as done.[79, 81–86] Subsequently, immunoblot analyses were performed using specific antibodies (see source companies, catalog numbers, and dilution ratios listed in **Table S3**). The specific protein bands were visualized by using Western Blotting Kit (Pierce, 35050).

### Mouse tumor xenograft models

Five-week-old female NOD-SCID mice were subcutaneously (sc) injected with H1792 or H2009 cells (1×10^6^) mixed in Matrigel solution (50%) into the hind flanks as previously done by us.[39, 40, 68, 72, 73] Tumor size was measured 2 to 3 times per week using a vernier caliper. When the tumor size reached 1000 mm^3, the mice were euthanized, and the tumors were excised for weight measurement and histological analysis. Throughout the study, all animals were maintained according to the University of Virginia (protocol#4372-11-21) and Richmond VAMC (IRBnet ID#1654460-4) approved IACUC protocols.

### Transcriptomics Analysis

The sequencing depth was 90 million reads per RNA sample from the two included cell lines with three replicates per cell line. The sequencing data was filtered with SOAPnuke (v1.5.6) (removing reads containing sequencing adapter and low-quality base ratio), then the clean reads were stored in FASTQ format. [87] Differential expression gene (DEG) analysis was performed using the DESeq2 with Q value ≤ 0.05. [63] KEGG/GO enrichment analysis of annotated different expression gene was performed by R Phyper (v120.0) based on Hypergeometric test. The significant levels of terms and pathways were corrected by Q value with a rigorous threshold (Q value ≤ 0.05).

### Statistical analysis

Graphing and statistics were performed using Prism GraphPad (Prism Software, San Diego, CA). The following models were used for statistical analysis as described, unpaired t-test with Welch’s correction were used when only two experimental groups were being compared. Repeated measures ANOVA were used when analyzing comparison groups that contained a dependent variable that was measured several times over the course of the experiment. One-way ANOVA with Tukey’s multiple comparisons test was used when comparing three or more independent groups. Two-way ANOVA with Šídák’s multiple comparisons were used when an examination of how the combination of two independent variables affected a dependent variable. All data reported as boxplot or mean (SD) where applicable; p < 0.05 was considered statistically significant. All bioinformatic analyses were conducted using the R program.

## Data and Materials Availability

All RNAseq data are in the process of submission to NCBI Sequence Read Archive (SRA) database for public access (Submission number: sub14341995). The GEO accession number is pending a database instrument update.

## Acknowledgements

The contents of this manuscript do not represent the views of the Department of Veterans Affairs or the United States Government. This work was mainly supported by the Veteran’s Administration (VA Merit Review, BX001792 (to CEC), VA Merit Review award, BX 006063 (C.E.C.), and a Senior Research Career Scientist Award, IK6BX004603 (to CEC)). This work was peripherally supported by way of research cores, methodology and technology development, and software development by the National Institutes of Health by way of P01 CA171983 (Project 1 to CEC), NIH/NCI Cancer Center Support Grant P30 CA044579, and the R01s AI139072 (to CEC), and DK126444 (to CEC). This work was also supported by funds from the Department of Medicine in the University of Virginia-School of Medicine in regard to software licensing and equipment purchases.

## Author Contributions

XX – designing research studies, conducting experiments, acquiring data, analyzing data, and writing the manuscript. HPM – developing and analyzing short and long read deep RNA sequencing data, manuscript method writing, and figure composition. ALL – conducting experiments, acquiring data, and analyzing data. CEC – overall supervision, designing research studies, analyzing data, providing reagents and funding, editing manuscript, and writing the manuscript.

