## Supplemental Data for "RNA splicing variants of the novel long non-coding RNA, CyKILR, possess divergent biological functions in non-small cell lung cancer"

#### Figure S1

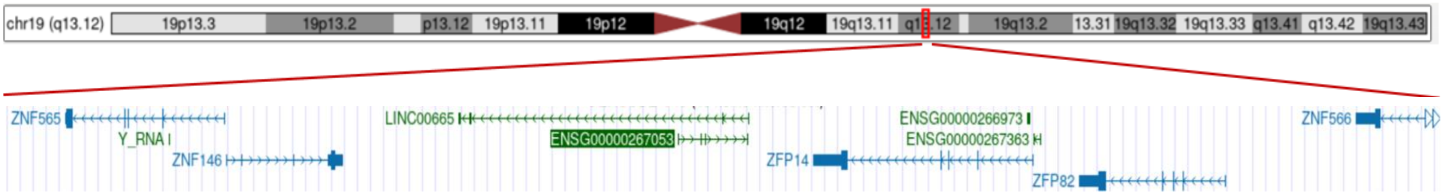


**Figure S1. The CyKILR locus (chromosome 19p13.12) and neighboring genes.** Genes located near the CyKILR transcriptional locus include *ZNF565*, *ZNF146*, *ZFP14*, *ZFP82*, and *ZNF566*.

### Figure S2

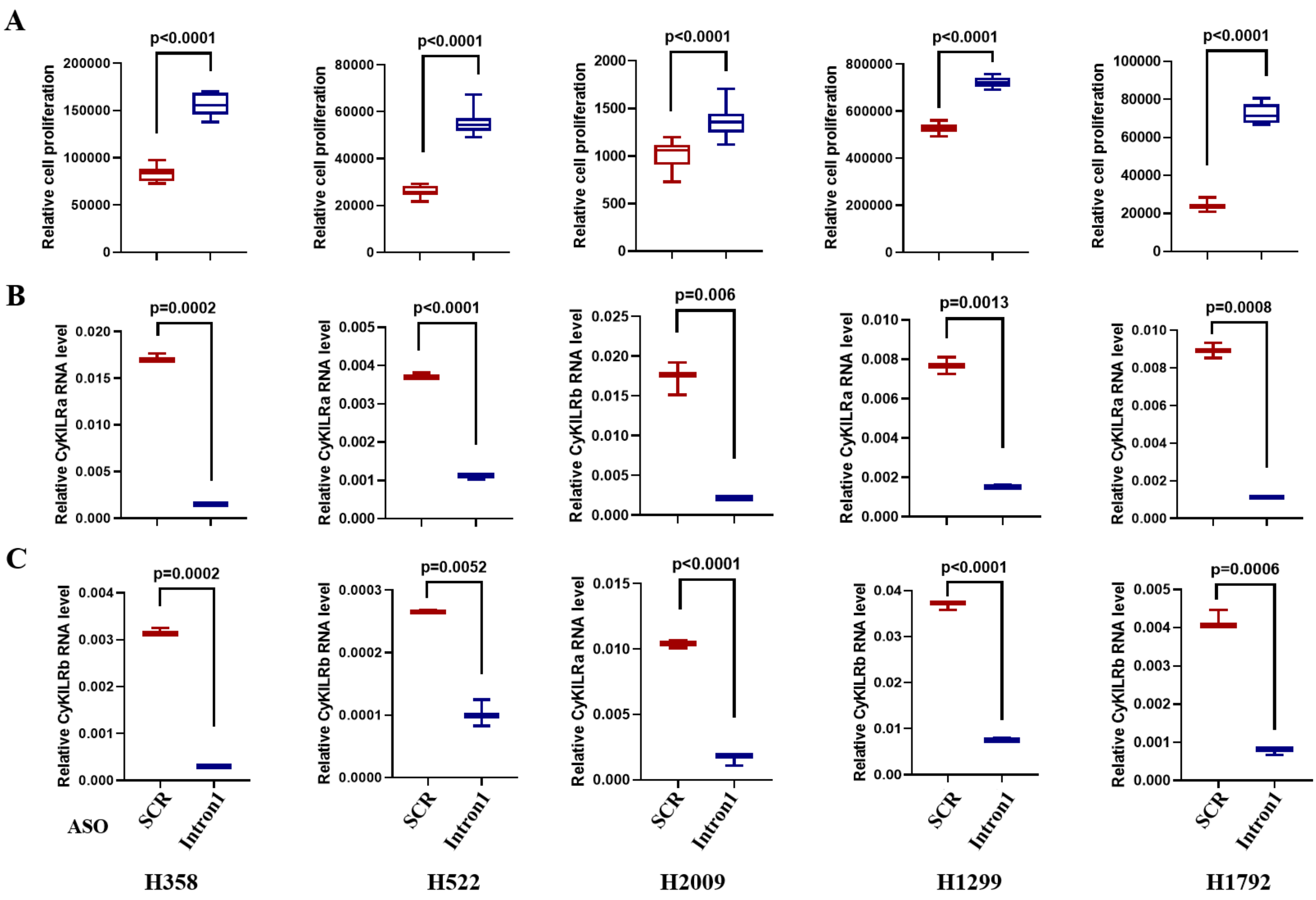


**Figure S2**. **The effect of depleting nascent CyKILR in H358, H522, H2009, H1299, and H1792 cell lines.** ASOs were designed to target the intronic region immediately downstream of exon 1 of CyKILR. Cells were transfected with either scrambled control ASO (SCR) or CyKILR Intron1 ASO (Intron1). Forty-eight hours post-transfection, the treated cells were subjected to either cell proliferation assays or qRT-PCR analysis for CyKILRa or b. **A.** CyKILR Intron1 ASO treatment enhanced cell proliferation across all five cell lines. **B.** CyKILR Intron1 ASO treatment reduced CyKILRa expression across all five cell lines. **C.** CyKILR Intron1 ASO treatment consistently reduced CyKILRb expression in the five cell lines. The data are presented as box plots (n ≥ 3 replicates), and statistical analysis was performed using the unpaired t-test with Welch's correction.

### Figure S3

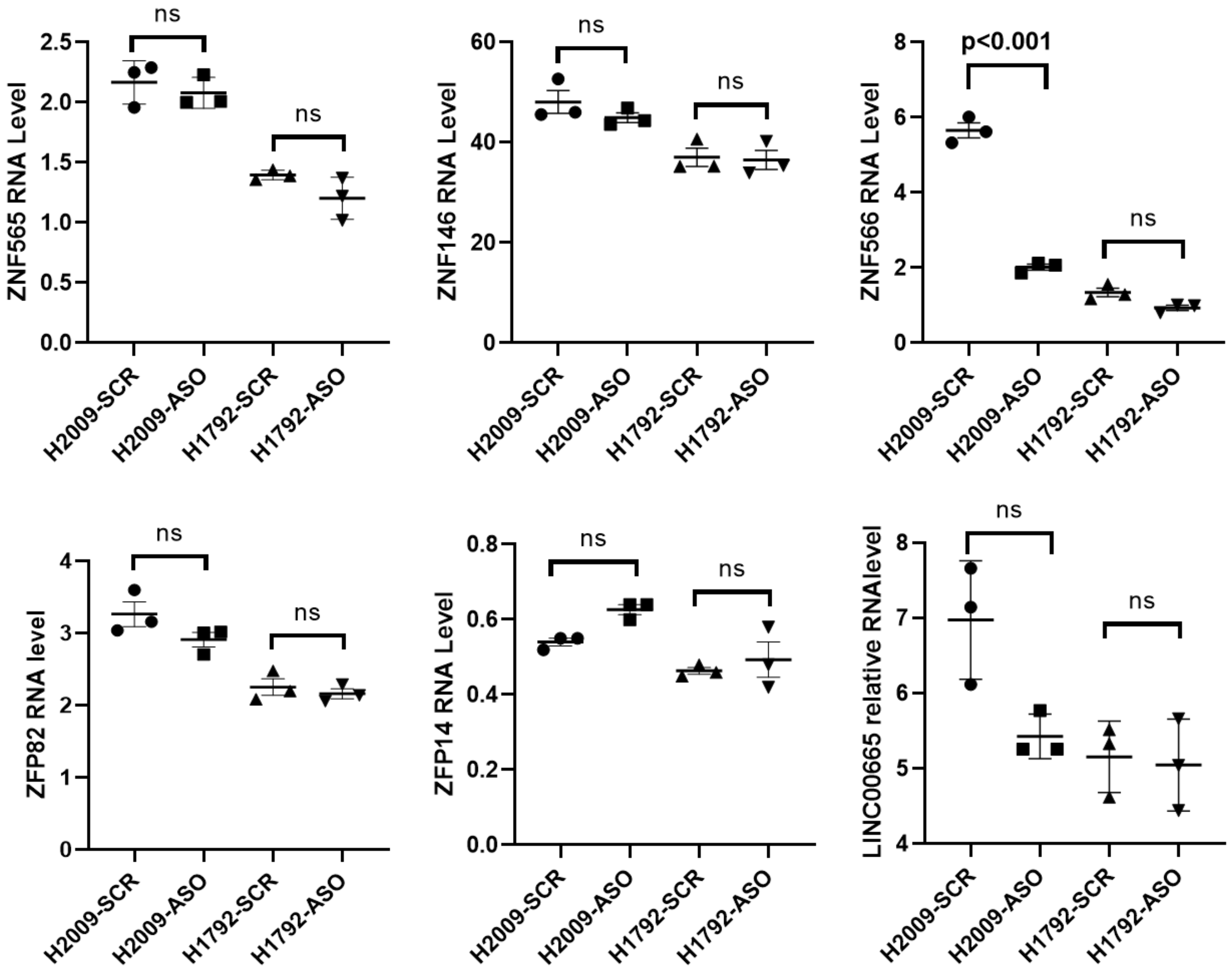


**Figure S3. Downregulation of CyKILRa does not affect the expression of adjacent/close proximity genes.** H2009 and H1792 cells were treated with either scrambled control ASO (SCR) or CyKILRa ASO (ASO), total RNA extracted, and then RNA was subjected to deep RNA sequencing. After filtering the data and aligning the clean reads, differential gene expression (DGE) analysis was conducted using the R package DESeq2. The expression of genes adjacent to CyKILR showed no significant changes when comparing the CyKILRa ASO-treated cells to the SCR-treated cells. The data are presented as dot plots (n= 3 replicates), and statistical analysis was performed using an unpaired t-test with Welch's correction.

### Figure S4

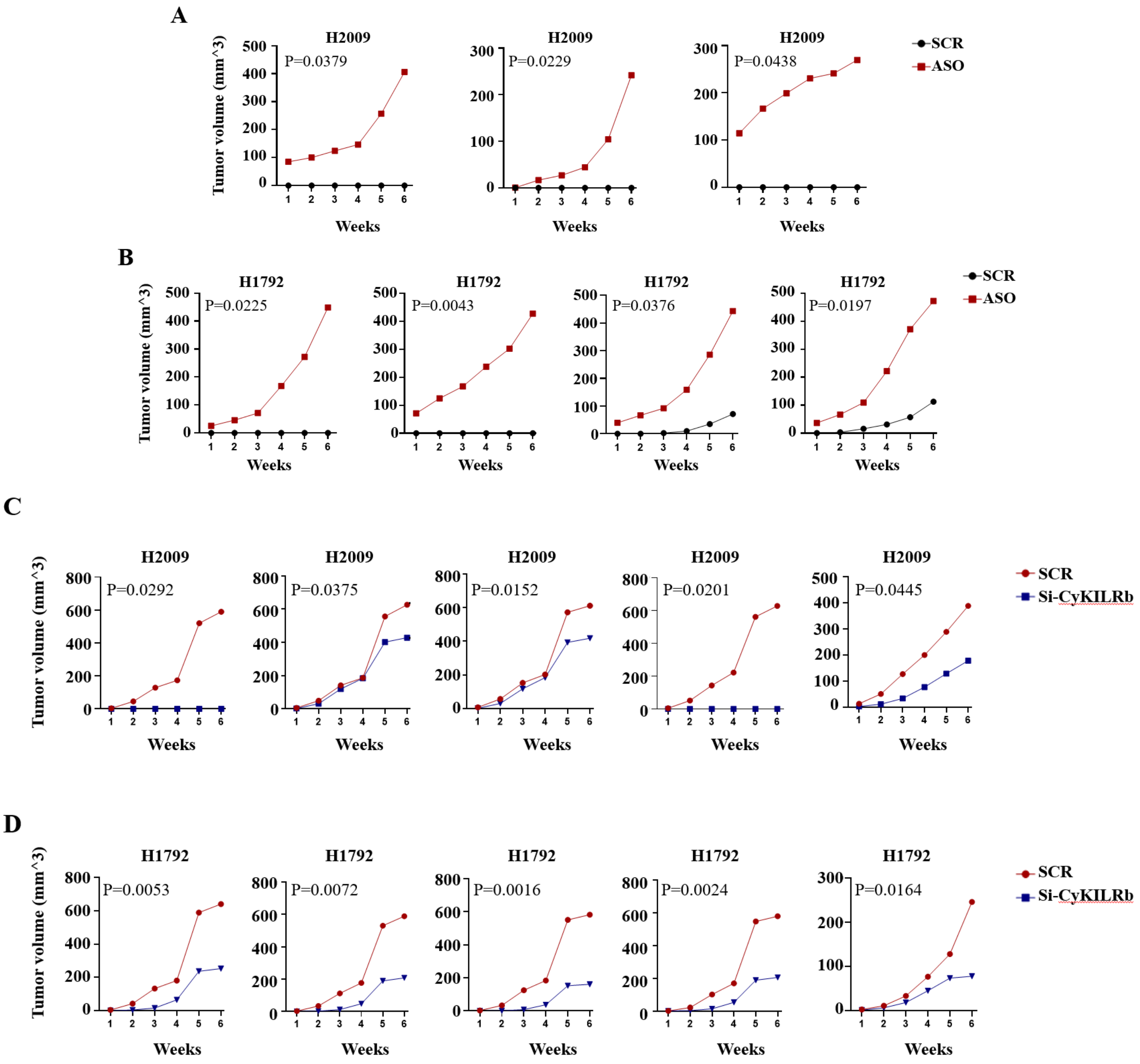


**Figure S4. Modulation of CyKILR splice variants demonstrates opposing effects on tumor growth in individual mice.** H2009 or H1792 cells were transfected with either (**A,B**) scrambled control ASO (SCR), CyKILRa ASO (ASO), (**C,D)**) scrambled control siRNA (SCR), or CyKILRb siRNA. After 48 hours post-transfection, the treated cells were mixed with Matrigel and injected subcutaneously (sc) into the flanks of five-week-old female NOD-SCID mice (1 x $10^6$ cells per mouse per flank). The left flank received Control-treated cells, and the right flank received CyKILRa ASO or CyKILRb siRNA knockdown cells. **A.** Tumor volume of SCR ASO or CyKILRa ASO treated H2009 for each depicted mouse. **B.** Tumor volume of SCR ASO or CyKILRa ASO treated H1792 for each depicted mouse. **C.** Tumor volume of SCR siRNA or CyKILRb siRNA treated H2009 for each depicted mouse. **D.** Tumor volume of SCR siRNA or CyKILRb siRNA treated H1792 for each depicted mouse. *p* values were assessed using Student t-tests with Welch's correction at the 6wk time point.

### Figure S5

**A.**

**H2009 ASO/SCR**

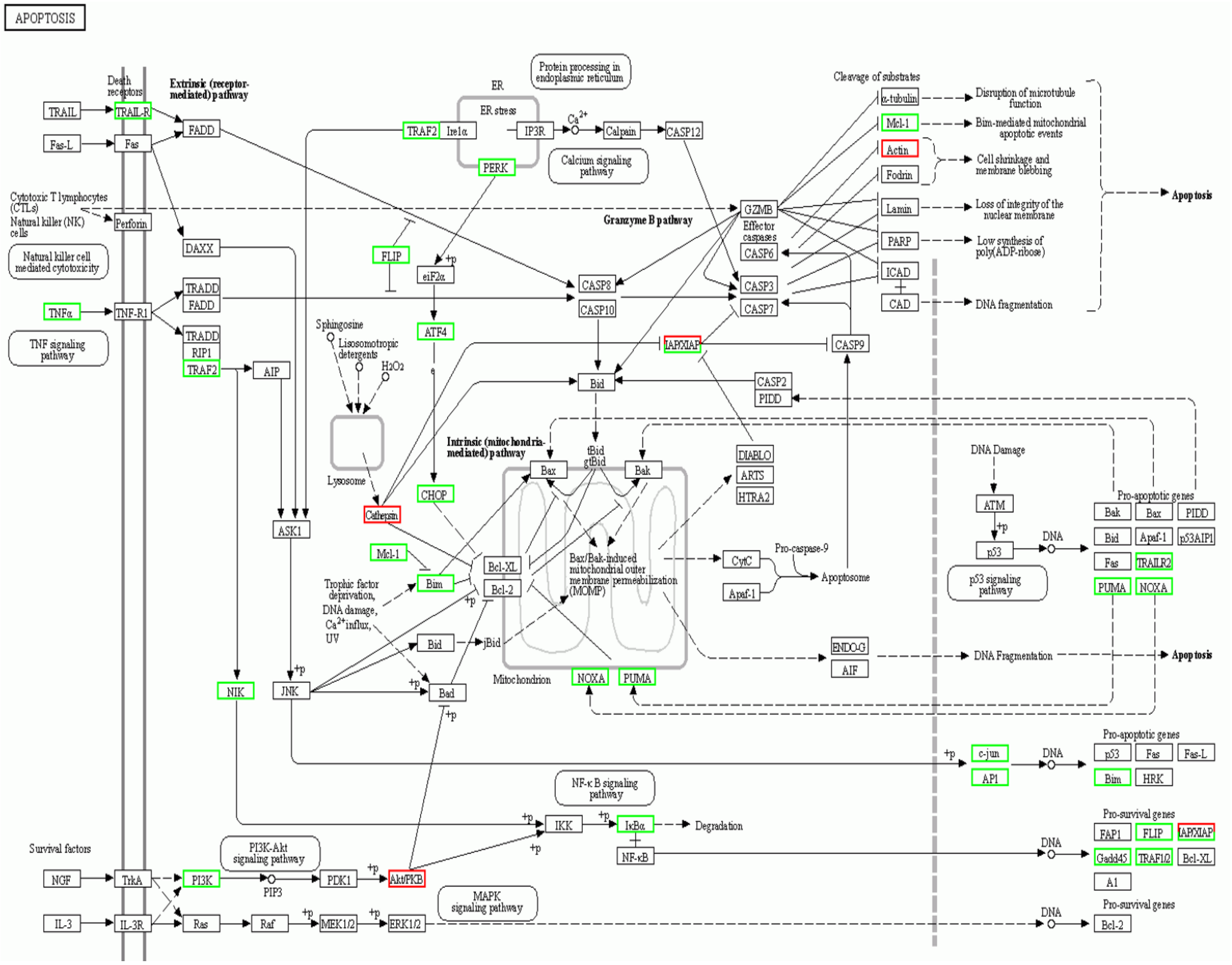

**B.**

**H1792 ASO/SCR**

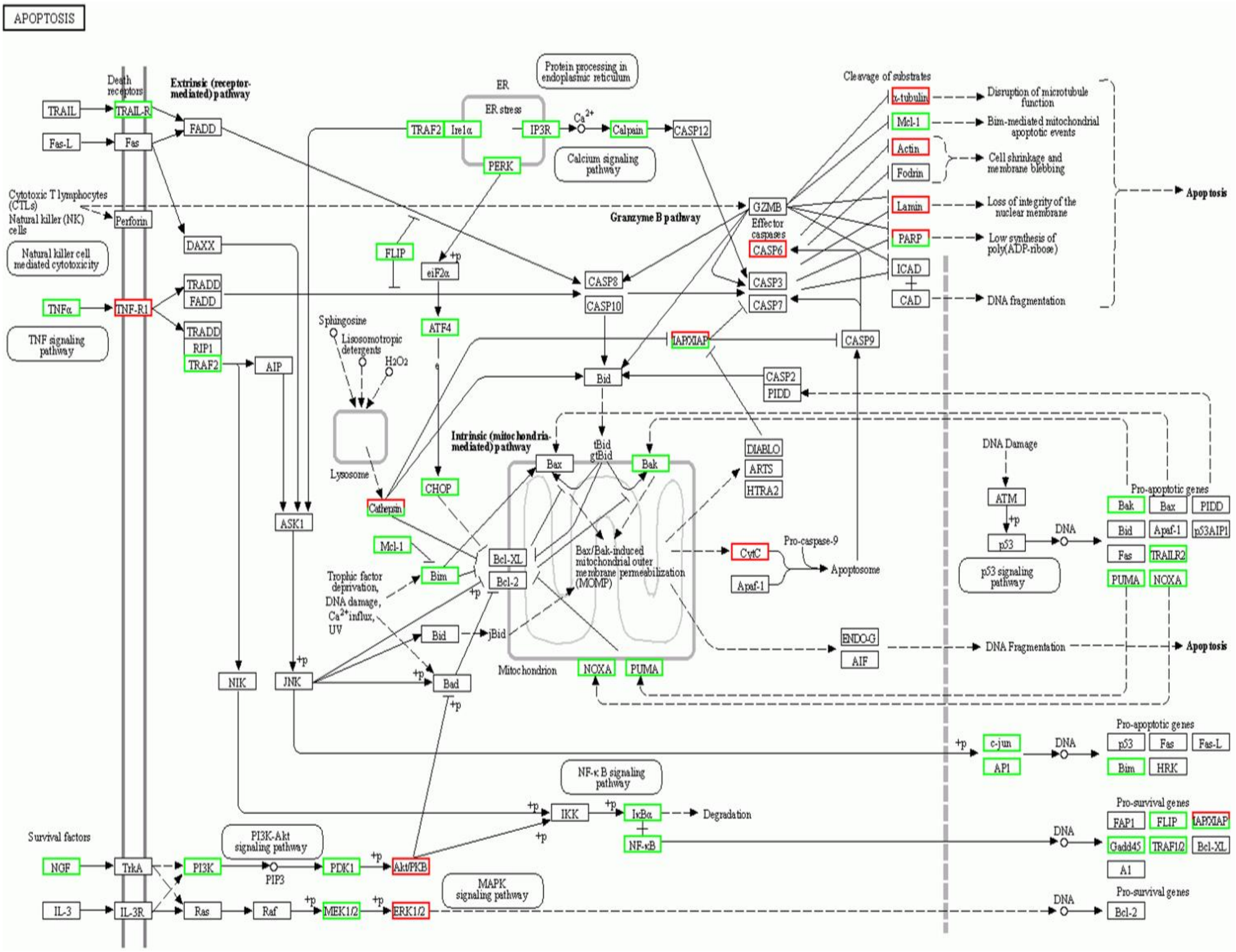


**Figure S5. CyKILRa regulates the apoptosis signaling pathway.** H2009 or H1792 cells treated with either scrambled control ASO (SCR) or CyKILRa ASO (ASO), total RNA extracted, and RNA underwent deep RNA sequencing. Following data filtering, clean-read alignment, gene expression, and differential expression gene analysis, KEGG enrichment analysis of annotated differentially expressed genes was performed using R Phyper. The ASO-treated cells group exhibited down-regulation of cell apoptosis pathways compared to the control-treated cells group. **A**. KEGG pathway of apoptosis, ASO versus SCR in H2009. **B**. KEGG pathway of apoptosis ASO versus SCR in H1792. Note: Green: down-regulation; Red: up-regulation.

#### Figure S6

**A.**

**H2009 ASO/SCR**

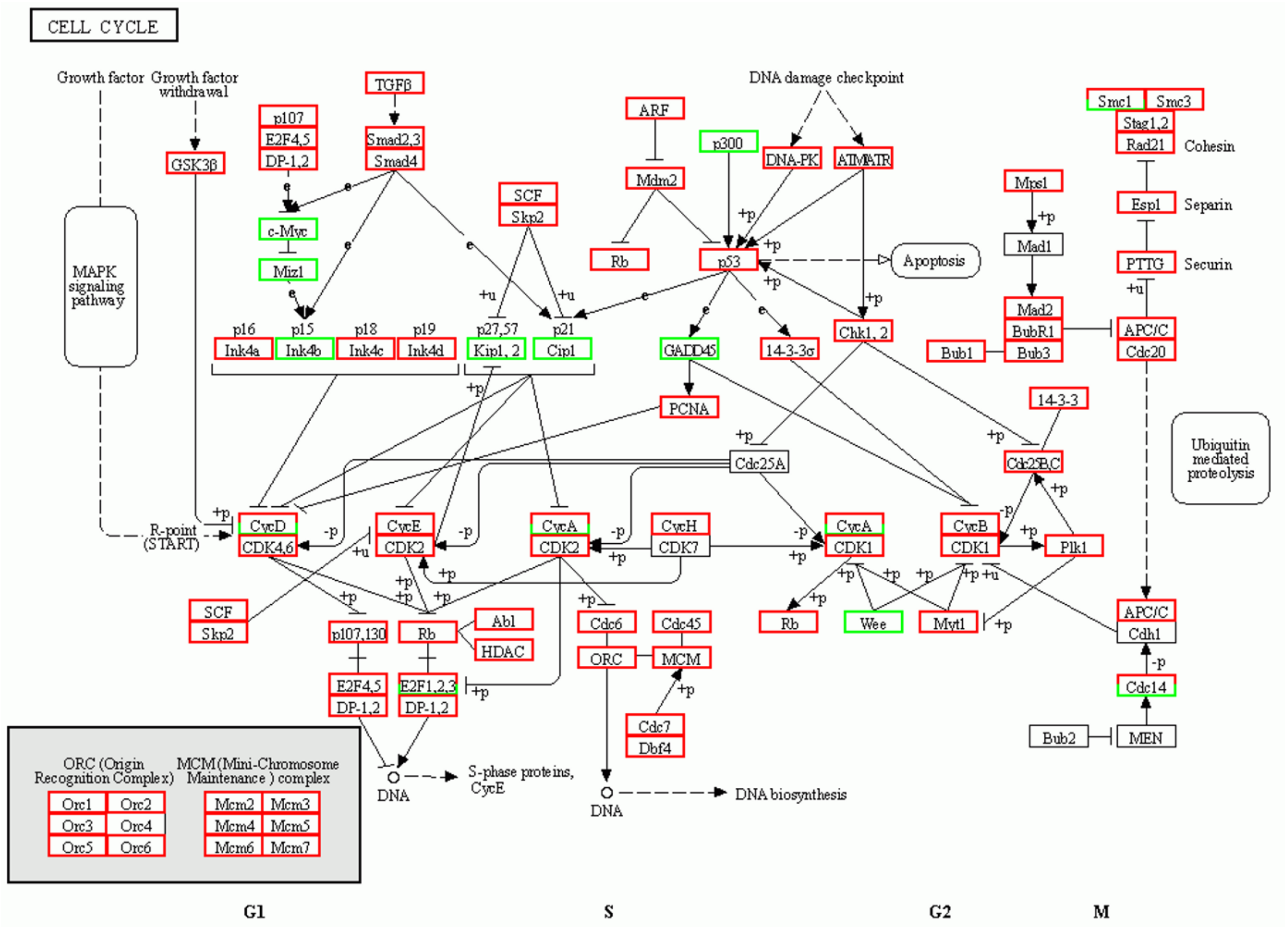

**B.**

**H1792 ASO/SCR**

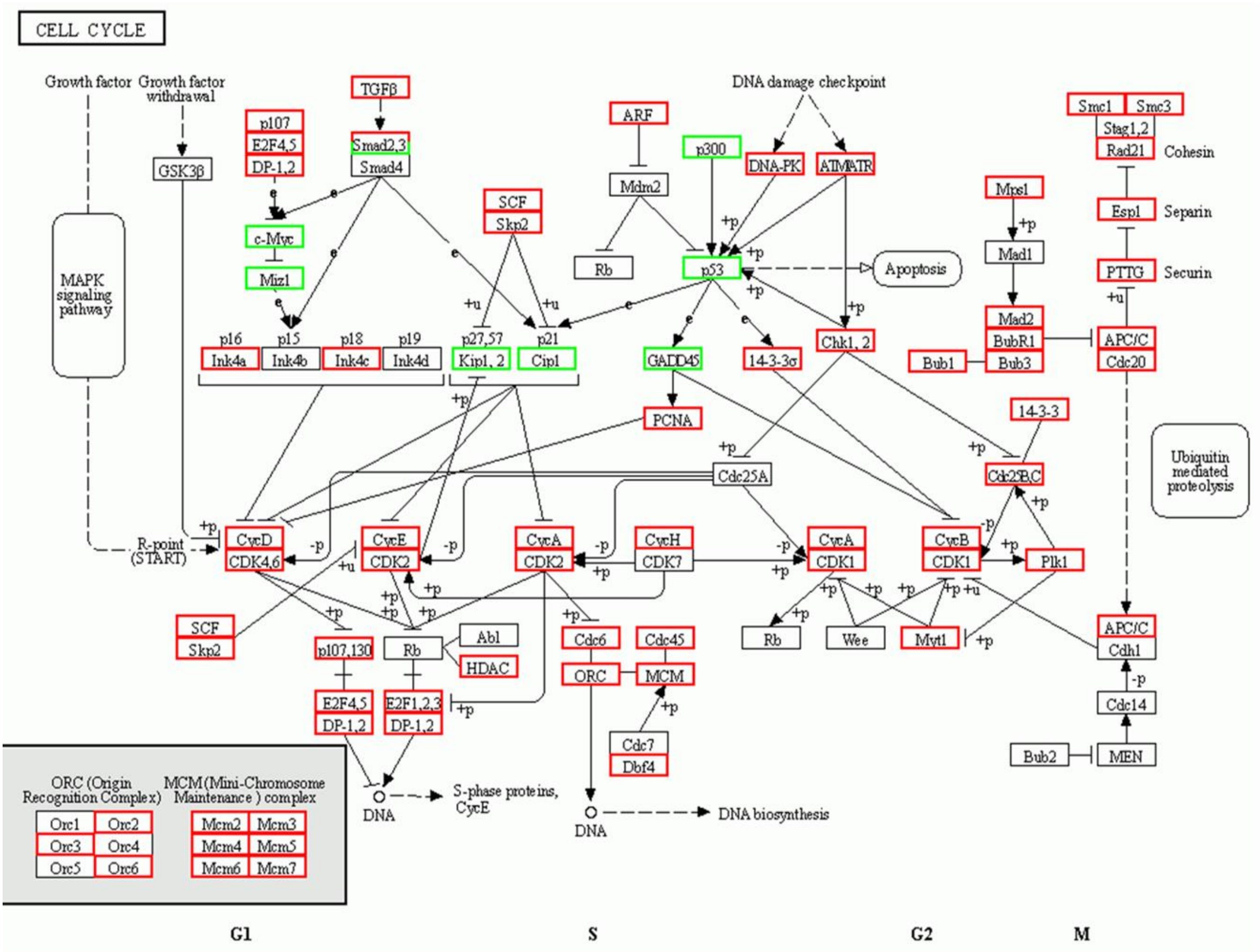


**Figure S6. CyKILRa regulates the cell cycle signaling pathway.** H2009 or H1792 cells treated with either scrambled control ASO (SCR) or CyKILRa ASO (ASO), total RNA extracted, and RNA underwent deep RNA sequencing. Following data filtering, clean-read alignment, gene expression, and differential expression gene analysis, KEGG enrichment analysis of annotated differentially expressed genes was performed using R Phyper. The ASO-treated cells group exhibited up-regulation of cell cycle pathways compared to the control-treated cells group. **A**. KEGG pathway of cell cycle, ASO versus SCR in H2009. **B**. KEGG pathway of cell cycle, ASO versus SCR in H1792. Note: Green: down-regulation; Red: up-regulation.

**Figure S7**

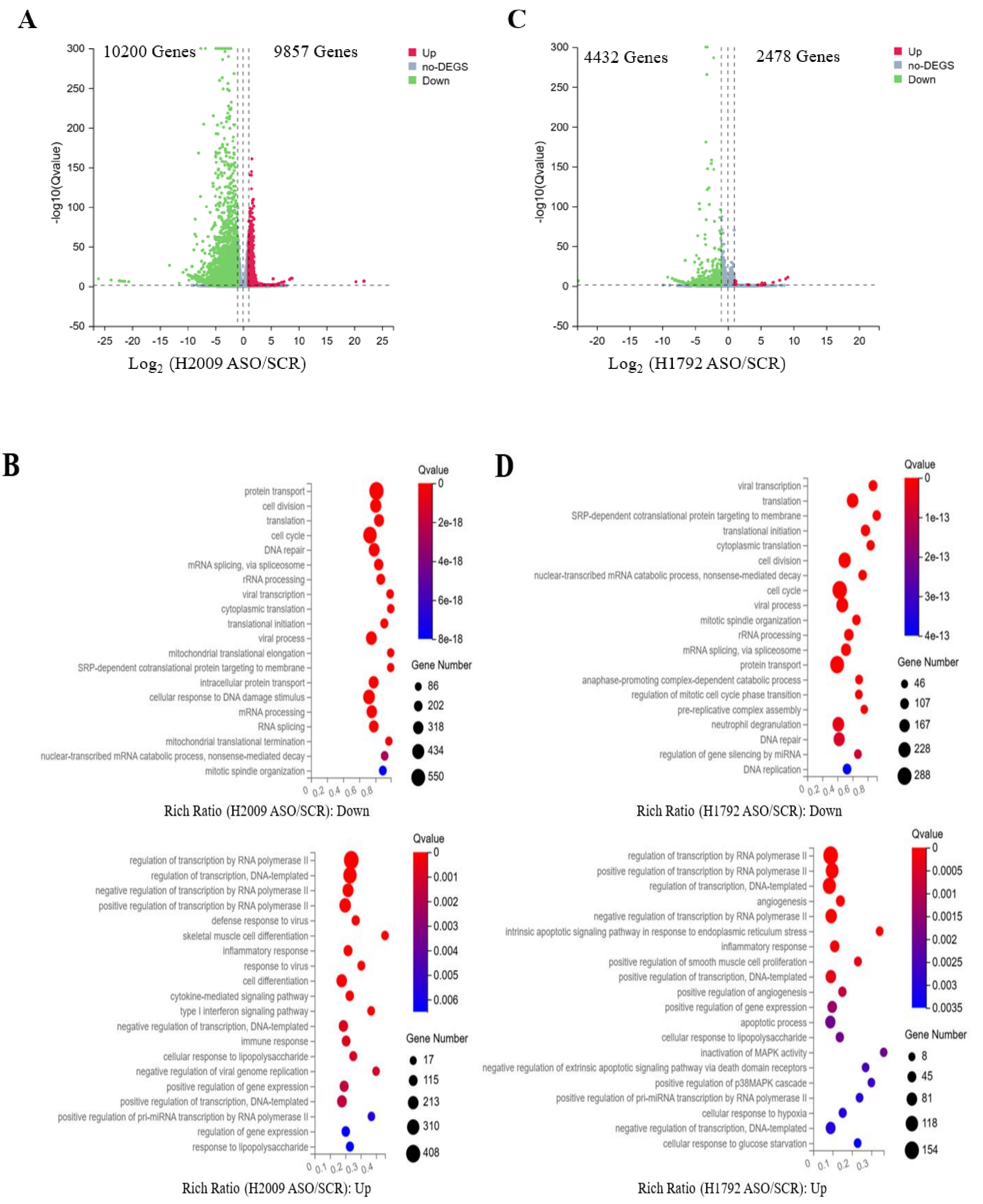
A
10200 Genes
9857 Genes
Up
no-DEGS
Down
-log10(Qvalue)
Log2 (H2009 ASO/SCR)
C
4432 Genes
2478 Genes
Up
no-DEGS
Down
-log10(Qvalue)
Log2 (H1792 ASO/SCR)
B
Qvalue
Gene Number
protein transport
cell division
translation
cell cycle
DNA repair
mRNA splicing, via spliceosome
rRNA processing
viral transcription
cytoplasmic translation
translational initiation
viral process
mitochondrial translational elongation
SRP-dependent cotranslational protein targeting to membrane
intracellular protein transport
cellular response to DNA damage stimulus
mRNA processing
RNA splicing
mitochondrial translational termination
nuclear-transcribed mRNA catabolic process, nonsense-mediated decay
mitotic spindle organization
Rich Ratio (H2009 ASO/SCR): Down
regulation of transcription by RNA polymerase II
regulation of transcription, DNA-templated
negative regulation of transcription by RNA polymerase II
positive regulation of transcription by RNA polymerase II
defense response to virus
skeletal muscle cell differentiation
inflammatory response
response to virus
cell differentiation
cytokine-mediated signaling pathway
type I interferon signaling pathway
negative regulation of transcription, DNA-templated
immune response
cellular response to lipopolysaccharide
negative regulation of viral genome replication
positive regulation of gene expression
positive regulation of transcription, DNA-templated
positive regulation of pri-miRNA transcription by RNA polymerase II
regulation of gene expression
response to lipopolysaccharide
Rich Ratio (H2009 ASO/SCR): Up
D
viral transcription
translation
SRP-dependent cotranslational protein targeting to membrane
translational initiation
cytoplasmic translation
cell division
nuclear-transcribed mRNA catabolic process, nonsense-mediated decay
cell cycle
viral process
mitotic spindle organization
rRNA processing
mRNA splicing, via spliceosome
protein transport
anaphase-promoting complex-dependent catabolic process
regulation of mitotic cell cycle phase transition
pre-replicative complex assembly
neutrophil degranulation
DNA repair
regulation of gene silencing by miRNA
DNA replication
Rich Ratio (H1792 ASO/SCR): Down
regulation of transcription by RNA polymerase II
positive regulation of transcription by RNA polymerase II
regulation of transcription, DNA-templated
angiogenesis
negative regulation of transcription by RNA polymerase II
intrinsic apoptotic signaling pathway in response to endoplasmic reticulum stress
inflammatory response
positive regulation of smooth muscle cell proliferation
positive regulation of transcription, DNA-templated
positive regulation of angiogenesis
positive regulation of gene expression
apoptotic process
cellular response to lipopolysaccharide
inactivation of MAPK activity
negative regulation of extrinsic apoptotic signaling pathway via death domain receptors
positive regulation of p38MAPK cascade
positive regulation of pri-miRNA transcription by RNA polymerase II
cellular response to hypoxia
negative regulation of transcription, DNA-templated
cellular response to glucose starvation
Rich Ratio (H1792 ASO/SCR): Up

E

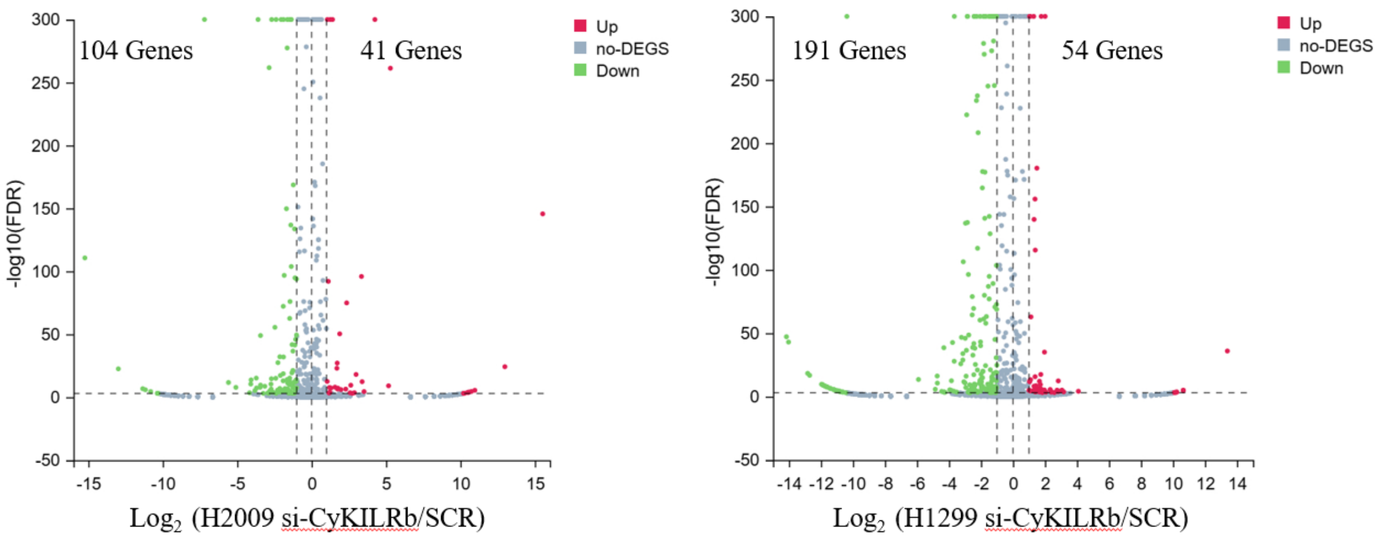


**Figure S7. Volcano plots and Gene Ontology (GO) analyses for CyKILRa and CyKILRb silencing.** H2009 or H1792 cells treated with either scrambled control ASO (SCR) or CyKILRa ASO (ASO), total RNA extracted, and RNA underwent deep RNA sequencing. H2009 or H1792 cells treated with either scrambled control siRNA (SCR) or CyKILRb siRNA (siCyKILRb), total RNA extracted, and RNA underwent small RNA sequencing. **A.** Volcano plot showing the number of down- and up-regulated genes under CyKILRa silencing in H2009. **B.** GO analysis of down- or up-regulated genes following CyKILRa knockdown in H2009. **C.** Volcano plot showing the number of down- and up-regulated genes under CyKILRa silencing in H1792. **D.** GO analysis of down- or up-regulated genes following CyKILRa knockdown in H1792. **E.** Volcano plot showing the number of down- and up-regulated genes under CyKILRb silencing in H2009 or H1299 cells.

### Figure S8

A

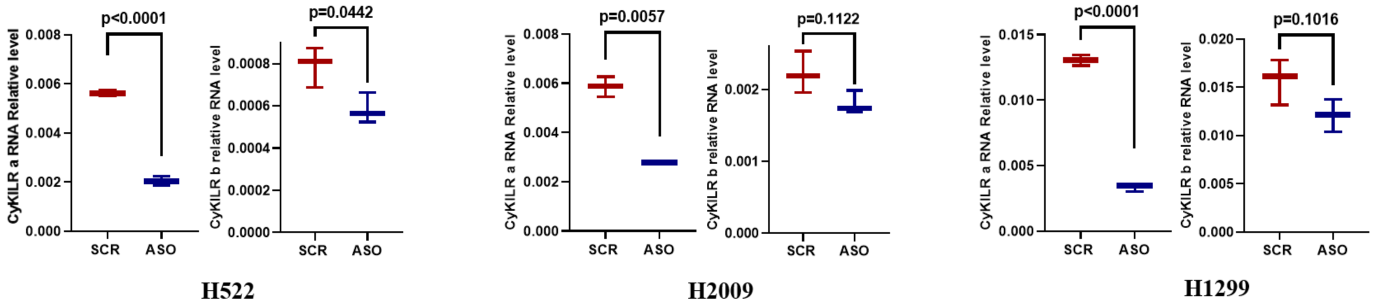


B

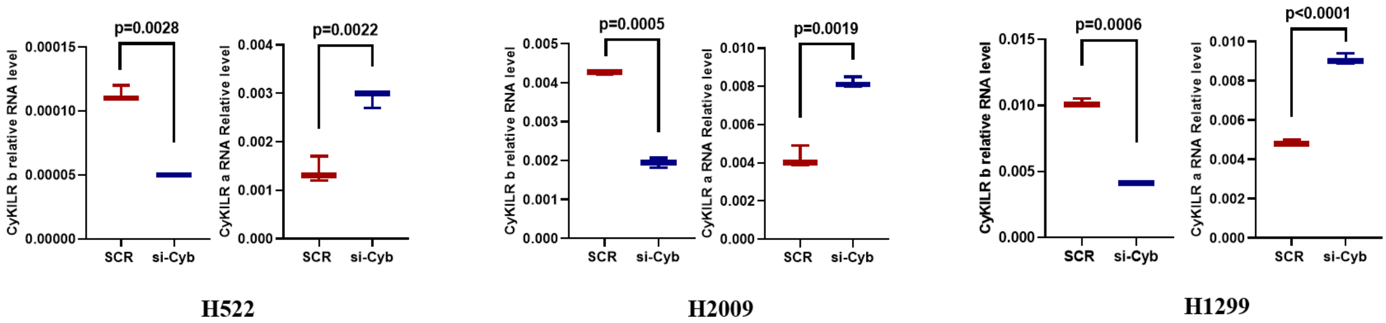


**Figure S8. Expression of each variant following the respective knockdown of CyKILRa and CyKILRb.** The depicted cell lines were treated with either scrambled control ASO (SCR), CyKILRa ASO (ASO), scrambled control siRNA (SCR), or CyKILRb siRNA (siCyKILRb), total RNA extracted, and RNA subjected to qRT-PCR for CyKILRa or b. **A.** CyKILRa knockdown had a minor effect on CyKILRb expression across three cell lines. **B.** CyKILRb knockdown significantly enhanced CyKILRa levels in all three cell lines. The data are presented as box plots ($n \geq 3$ replicates), and statistical analysis was performed using an unpaired t-test with Welch's correction.

**Table S1. Top differentially expressed long non-coding RNAs (lncRNAs) identified in a comparison between CDKN2A/LKB1 wild-type and mutant non-small cell lung cancer (NSCLC) cell lines.** Two cell lines from each group were used in the analysis: H1299 and H1792 (wild-type) and H460 and A549 (mutant). Gene expression data were analysed and ranked using the DESeq2 pipeline. The table columns include Base Mean, representing the average expression of each gene across all samples; log2 Fold Change, reflecting the change in gene expression between mutant and wild-type groups (with positive values indicating upregulation in the mutant group); lfc SE, the standard error of the log2 fold change; and stat, the Wald test statistic. P-values were calculated using the Wald test and adjusted for multiple comparisons using the Benjamini-Hochberg method (p-adj).

| ENSG ID | Base Mean | log2 Fold Change | lfc SE | stat | p-value | p-adj |
|---|---|---|---|---|---|---|
| **ENSG00000267053** | 1505.14 | 9.04 | 0.73 | 12.37 | 3.60E-35 | 2.54E-33 |
| ENSG00000233854 | 114.17 | 8.41 | 0.75 | 11.26 | 2.11E-29 | 1.16E-27 |
| ENSG00000234286 | 47.70 | 8.38 | 0.79 | 10.61 | 2.75E-26 | 1.25E-24 |
| ENSG00000266560 | 102.55 | 9.56 | 0.92 | 10.38 | 3.08E-25 | 1.32E-23 |
| ENSG00000289502 | 53.30 | 8.95 | 0.87 | 10.30 | 6.77E-25 | 2.85E-23 |
| ENSG00000267886 | 67.23 | 8.46 | 0.85 | 9.95 | 2.64E-23 | 1.02E-21 |
| ENSG00000250284 | 196.09 | 9.56 | 1.01 | 9.45 | 3.33E-21 | 1.11E-19 |
| ENSG00000275874 | 74.72 | 9.03 | 1.00 | 9.04 | 1.64E-19 | 4.83E-18 |
| ENSG00000257056 | 40.22 | 8.54 | 0.95 | 9.01 | 2.10E-19 | 6.15E-18 |

**Table S2. Sequence of RT-PCR primers, siRNA, and ASO**

| Gene Target | RT-qPCR | |
|---|---|---|
| CDKN2A | Assay ID: Hs00923894_m1(ThermoFisherScientific, Cat: 4331182) | |
| *STK11*/LKB1 | Assay ID: Hs00975986_m1(ThermoFisherScientific, Cat: 4331182) | |
| GAPDH | Assay ID: Hs02758991_g1 (ThermoFisherScientific, Cat: 4331182) | |
| CyKILR (Total) | Forward (E5) | 5’-TTCTCCTCTGCAACCCAATG-3’ (Sense) |
| | Reverse (E5) | 5’-GTTTCCTTCAAGCCGTACCT-3’ (AntiSense) |
| | Probe (E5) | 5’-TTTCCAGAATCCTTTGGCGGTCCC-3’ (AntiSense) |
| CyKILRa | Forward (E3) | 5’-CAGGGAGTGAAGCCTTTGT-3’ (Sense) |
| | Reverse (E3) | 5’-GCACCTCCAGGCCATTT-3’ (Antisense) |
| | Probe (E3) | 5’-CCCAAGCAACAGAGTGAGGGTCTG-3’ (Sense) |
| CyKILRb | Forward (E1) | 5’-GTCCAGCAGCAGGAAGAAA-3’ (Sense) |
| | Reverse (E5) | 5’-GCTCCTAAGCTTGGTTGTGT-3’ (Antisense) |
| | Probe (E1-E5) | 5’-AAGAAACCTCAGACAGATCGCCGG-3’ (Sense) |
| | **(Multiplex) RT-PCR** | |
| CyKILRa | Forward (E3) | 5’-CACTGTTAGGGTCGGGGATG-3’ |
| | Reverse (E3) | 5’-TGCAGTCTGTGAGTCCCCTA-3’ |
| CyKILRb | Forward (E1) | 5’-GTCCAGCAGCAGGAAGAAA-3’ |
| | Reverse (E1-E5) | 5’-GCTTGGTTGTGTGCGCCGCA-3’ |
| | **siRNA** | |
| CyKILRb | UGC GGC GCA CAC AAC CAA GC UU | |
| | GCG CGG ACG GUG UGC GGC G UU | |
| | GGA CGG UGU GUG CGG CGC ACA CUU | |
| Scramble (siRNA) | Silencer Select Negative Control No.1 siRNA (ThermoFisherScientific, Cat: 4390844) | |
| | **Antisense Oligonucleotides (ASO)** | |
| CyKILRa | 1 | mG*mC*mU*mC*mC*A*G*A*G*A*C*C*C*A*C*mU*mG*mU*mU*mA (sense)<br>mU*mA*mA*mC*mA*G*T*G*G*G*T*C*T*C*T*mG*mG*mA*mG*mC (Antisense) |
| | 2 | mC*mC*mU*mC*mU*A*G*C*T*C*A*G*C*C*C*mC*mA*mA*mG*mC (sense)<br>mG*mC*mU*mU*mG*G*G*G*C*T*G*A*G*C*T*mA*mG*mA*mG*mG (Antisense) |
| | 3 | mA*mG*mU*mA*mA*C*A*G*A*G*G*G*A*C*A*mC*mG*mC*mA*mG (sense)<br>mC*mU*mG*mC*mG*T*G*T*C*C*C*T*C*T*G*mU*mU*mA*mC*mU (Antisense) |
| Scramble (ASO) | | mA*mG*mG*mA*mT*T*A*G*C*G*A*T*A*A*G*mT*mC*mG*mA*mA(sense)<br>mU*mU*mC*mG*mA*C*U*U*A*U*C*G*C*U*A*A*mU*mC*mC*mU(Antisense) |
| CyKILR intron1 | 1 | mA*mG*mU*mG*mG*A*G*C*C*C*T*T*G*T*G*mA*mU*mG*mA*mC (sense)<br>mG*mU*mC*mA*mU*mC*A*C*A*A*G*G*G*mC*T*C*mC*mA*mC*mU(Antisense) |
| | 2 | mC*mU*mU*mG*mU*G*A*T*G*A*C*G*A*G*C*mA*mC*mA*mG*mG(sense)<br>mC*mC*mU*mG*mU*G*C*T*C*G*T*C*A*T*C*mA*mC*mA*mA*mG(Antisense) |

**Table S3. Antibodies for Western immunoblotting**

| Antibody | Company | Catalog number | Dilution |
|---|---|---|---|
| CDKN2A/p14ARF | Abcam | ab185620 | 1:1000 |
| P16-INK4A | Proteintech | 10883-1-AP | 1:1000 |
| LKB1 | ThermoFisherScientific | AHO1392 | 1:500 |
| β-actin | ThermoFisherScientific | MA1-140 | 1:10000 |
